# Barth syndrome cellular models have dysregulated respiratory chain complex I and mitochondrial quality control due to abnormal cardiolipin

**DOI:** 10.1101/2021.01.06.425502

**Authors:** Arianna F. Anzmann, Olivia L. Sniezek, Alexandra Pado, Veronica Busa, Frédéric Maxime Vaz, Simion D. Kreimer, Robert Norman Cole, Anne Le, Brian James Kirsch, Steven M. Claypool, Hilary J. Vernon

**Author notes:** **Corresponding author at:** Department of Genetic Medicine, Johns Hopkins University School of Medicine, 733 N Broadway, MRB 512 Baltimore, Maryland, USA. (H.J. Vernon).

## Abstract

Barth syndrome (BTHS) is an X-linked genetic condition caused by defects in *TAZ*, which encodes a transacylase involved in the remodeling of the inner mitochondrial membrane phospholipid, cardiolipin (CL). As such, CL has been implicated in numerous mitochondrial functions, and the role of defective CL in the clinical pathology of BTHS is under intense investigation. We used untargeted proteomics, shotgun lipidomics, gene expression analysis, and targeted metabolomics to identify novel areas of mitochondrial dysfunction in a new model of TAZ deficiency in HEK293 cells. Functional annotation analysis of proteomics data revealed abnormal regulation of mitochondrial respiratory chain complex I (CI), driven by the reduced abundance of 6 CI associated proteins in TAZ-deficient HEK293 cells: MT-ND3, NDUFA5, NDUFAB1, NDUFB2, NDUFB4, and NDUFAF1. This resulted in reduced assembly and function of CI in TAZ-deficient HEK293 cells as well as BTHS patient derived lymphoblast cells. We also identified increased abundance of PARL, a rhomboid protein involved in the regulation of mitophagy and apoptosis, and abnormal downstream processing of PGAM5, another mediator of mitochondrial quality control, in TAZ-deficient cells. Lastly, we modulated CL via the phospholipase inhibitor bromoenol lactone and the CL targeted SS-peptide, SS-31, and showed that each is able to remediate abnormalities in CI abundance as well as PGAM5 processing. Thus, mitochondrial respiratory chain CI and PARL/PGAM5 regulated mitochondrial quality control, both of whose functions localize to the inner mitochondrial membrane, are dysregulated due to TAZ deficiency and are partially remediated via modulation of CL.

## Introduction

Barth syndrome (BTHS, MIM#302060) is a rare X-linked inborn error of mitochondrial phospholipid metabolism caused by variants in the gene TAFAZZIN (*TAZ*) (1–3). *TAZ* encodes a transacylase essential for the remodeling and maturation of the mitochondrial phospholipid cardiolipin (CL) (1, 4). CL, primarily localized to inner mitochondrial membrane, has many key functions, including roles in maintaining mitochondrial cristae structure, organization of respiratory complexes, protein import, fusion, fission, and cellular signaling (1). TAZ deficiency results in abnormal CL content, including an accumulation of the remodeling intermediate monolysocardiolipin (MLCL), decreased remodeled CL, and a shift towards saturated CL species (1, 5). An elevated MLCL:CL ratio is the pathognomonic metabolic defect in BTHS and is found in 100% of affected individuals (6).

BTHS is a multisystem disorder characterized by prenatal onset of left ventricular noncompaction, early onset cardiomyopathy, skeletal myopathy, growth abnormalities, and neutropenia among other features, and is the only known Mendelian disorder of CL metabolism (1, 7–9). Despite knowledge of the primary metabolic defect in BTHS, there is limited knowledge of downstream mechanisms of cellular pathogenesis, and consequently there is a dearth of targets for therapeutic intervention and clinical monitoring (10). In addition to BTHS, CL abnormalities have been described in common conditions such as idiopathic cardiomyopathy, fatty liver disease, and diabetes (1, 11–14). Consequently, studies in BTHS have the potential to illuminate pathophysiology in a range of common diseases.

To identify novel and unappreciated cellular pathways impacted by TAZ deficiency we employed a discovery-based approach in a new HEK293 TAZ-deficient cellular model, starting with untargeted proteomics followed by functional analysis, and validation of pathways of interest in both HEK293 TAZ-deficient cells and patient derived lymphoblastoid cell lines (LCLs). With this approach we characterized two major areas of dysfunction in the inner mitochondrial membrane: complex I (CI) of the mitochondrial respiratory chain and mitochondrial quality control (MQC).

Prior studies seeking to define the mitochondrial pathology of TAZ deficiency have described abnormal assembly and function of the mitochondrial respiratory complexes (15–17). In agreement with these findings, we identified aberrations in mitochondrial respiratory chain protein abundance and enzymatic function, specifically in CI. Significantly, our findings show that TAZ deficiency results in the reduced CI mRNA expression with evidence for distinct regulation in differing cell types.

In addition to expanding our current understanding of respiratory chain abnormalities in TAZ deficiency, we also identified novel abnormalities in regulators of MQC. We uncovered aberrant abundance of the MQC protein, mature mitochondrial PARL, which was accompanied by altered cleavage of the downstream MQC mediator and PARL target, PGAM5. PGAM5 cleavage abnormalities were further amplified by uncoupling of mitochondrial oxidative phosphorylation, suggesting that baseline MQC abnormalities in TAZ-deficient cells can be exacerbated with additional stress.

Finally, we found that modulating CL normalizes gene expression of CI subunits, normalizes the abundance of CI holoenzyme, and remediates the aberrant ratio of cleaved to full length PGAM5. Thus, abnormal CL in TAZ deficiency has a direct role in dysregulating both CI of the mitochondrial respiratory chain and MQC.

## Results

### Generation of a HEK293 TAZ-KO model, TAZ^Δ45^

We used CRISPR/Cas9 genome editing in HEK293s to create a novel TAZ-deficient cellular model. Using two single guide RNAs (sgRNAs) targeting exon 2 of *TAZ*, we isolated 3 individual clones with a resultant 45bp deletion at the 3’-end of exon 2; *TAZ*^Δ45.4^, *TAZ*^Δ45.5^, *TAZ*^Δ45.6^ (Fig.S1). The deletion encompasses a predicted acyltransferase domain and covers an area of *TAZ* where multiple pathological variants have been described, such as p.R57L and p.H69Q (Fig. S1) (3). The 45bp in-frame deletion is not predicted to result in NMD (Fig. 1A). However, the deletion resulted in undetectable expression of TAZ in all three clones (Fig. 1B). In the absence of TAZ there was no significant difference in the abundance of cytosolic and mitochondrial proteins/immunoblotting controls; GRP75, ß-actin, VDAC1, and TOM20 (Fig. S2).

**Figure 1.**
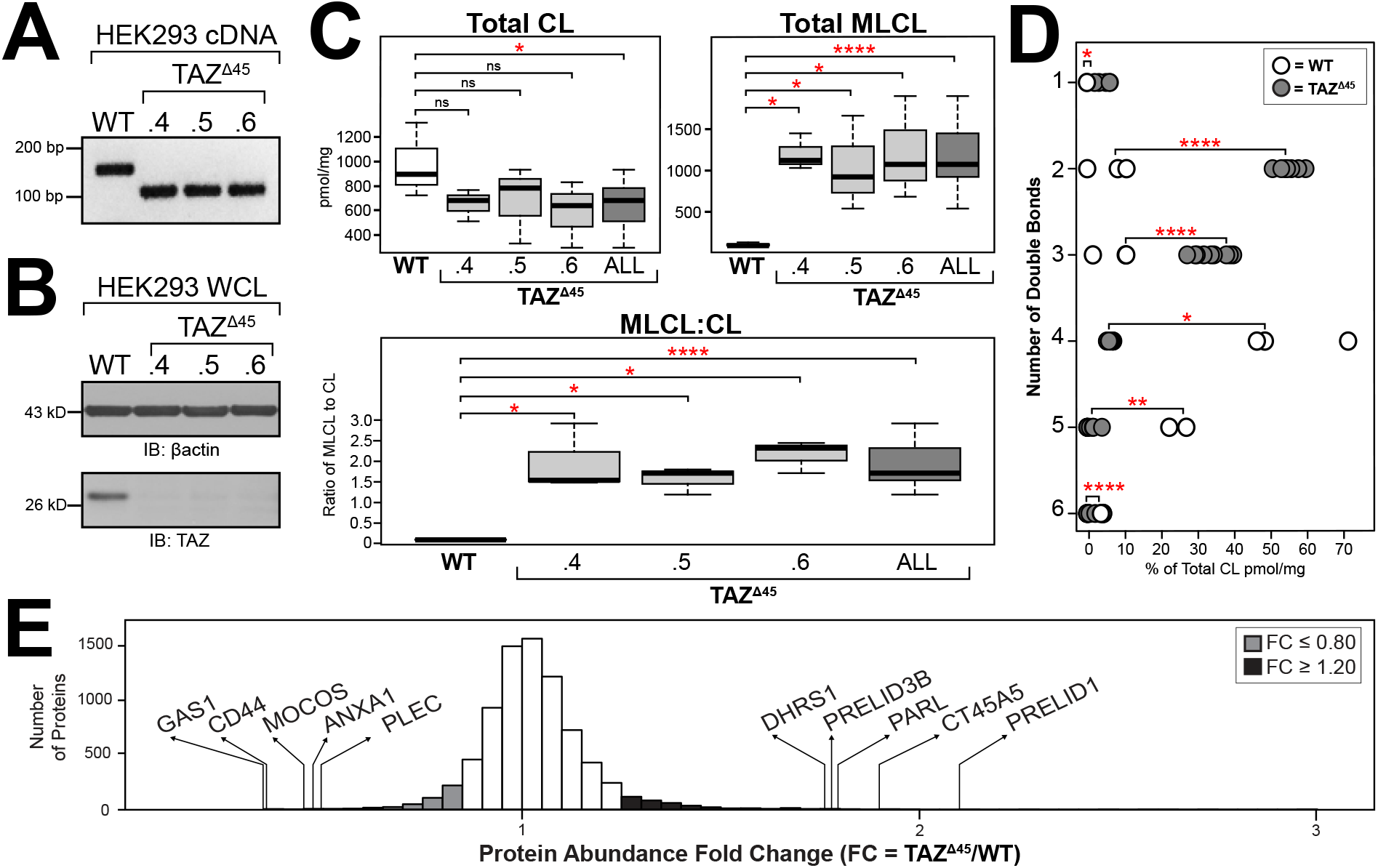
TAZD45 720 HEK293 genetic characterization, CL profiling and proteomics analysis. **(A)** RT-PCR of RNA extracted from the HEK293 *TAZ^Δ45^* clones using primers specific to the region of *TAZ* being edited. **(B)** Whole cells lysate (45ug) of the indicated lines immunoblotted for TAZ and loading control β-actin. **(C)** The abundance of CL and MLCL in WT (n=3) and *TAZ*^Δ45^ clones (n=3, per clone) was determined by shotgun lipidomics via mass spectrometry. **(D)** The distribution of double bonds across CL species was determined as a percentage of total CL. **(E)** 7713 proteins were identified and quantified with TMT 10-plex mass spectrometry of whole cell lysate (200ug) of *TAZ*^Δ45^ (n=3) and WT cells (n=3). Protein abundance fold change (FC) calculated by dividing the average abundance per protein identified in *TAZ*^Δ45^ cells by WT average abundance per protein. The genes that encode the most significantly reduced proteins in *TAZ*^Δ45^ cells are *GAS1* (FC=0.35, p=2.5 x 10-4), *CD44* (FC=0.367, p=6.6 x 10-4), *MOCOS* (FC=0.45, p=0.011), *ANAX1* (FC=0.478, p=0.01), and *PLEC* (FC=0.491, p=2.7 x 10-4). The genes that encode the most significantly increased proteins in *TAZ*^Δ45^ cells are *PRELID1* (FC=2.125, p=8.4 x 10-4), *CT45A5* (FC=1.909, p=0.023), *PARL* (FC=1.815, p=0.016), *PRELID3B* (FC=1.795, p=3.9 x 10-4), and *DHRS1* (FC=1.774, p=0.016). Proteins with a fold change (FC) ≤ 0.80 are highlighted in grey (n=215) and proteins with a FC ≥ 1.20 are highlighted in black (n=621). Significant differences are indicated; * ≤ 0.05, ** ≤ 0.005, *** ≤ 0.0005, **** ≤ 0.00005.

Shotgun lipidomics via mass spectrometry revealed a significant decrease in CL, a significant increase in MLCL, and a significantly increased MLCL:CL ratio (p=0.03, p=4.9 × 10^−5^, p=4.6 × 10^−6^, respectively) (Fig. 1C). TAZ-based remodeling is characterized by the incorporation of unsaturated acyl chains compared to nascent CL. Of the 31 CL species assessed, the *TAZ*^Δ45^ clones had a significant increase in CL containing 1 to 3 double bonds (p=0.007, p=1.5 × 10^−9^, p=7.4 × 10^−6^, respectively) and a significant decrease in CL containing 4 to 6 double bonds (p=0.03, p=0.003, p=4.2 × 10^−9^,, respectively), highlighting the loss of TAZ-based remodeling (Fig. 1D). Collectively, the *TAZ*^Δ45^ clones recapitulate the pathognomonic metabolic defect of BTHS and validate the *TAZ*^Δ45^ clones as TAZ-deficient cellular models of BTHS.

We were able to amplify and Sanger sequence 5 of the top 10 predicted off-target sites, 5 off-target sites per sgRNA, which revealed no detectable off-target CRISPR/Cas9 genome editing activity (Table S1). The other 5 off-target sequences were not able to be amplified likely due to highly repetitive sequences and/or increased GC base pair content. None of the top 10 predicted off-target sites were located in coding regions. In order to mitigate consequences of undetected off-target editing in any one of the clones, the clones, *TAZ*^Δ45.4^, *TAZ*^Δ45.5^, *TAZ*^Δ45.6^, were combined at a 1:1:1 ratio to create the cellular model *TAZ*^Δ45^.

### Differentially abundant proteins in *TAZ*^Δ45^ cells reveal downstream cellular dysfunction due to TAZ deficiency

Shotgun proteomics analysis identified a total of 7713 proteins in HEK293 WT and *TAZ*^Δ45^ cells. To focus our downstream workflow on proteins with differential abundance between the WT and *TAZ*^Δ45^ cells, we selected proteins with a protein abundance fold change (FC, *TAZ*^Δ45^/ WT) less than or equal to 0.80 (FC ≤ 0.80) and proteins with a FC greater than or equal to 1.2 (FC ≥ 1.20) (Fig. 1E). Based on these criteria, there were a total of 836 differentially abundant proteins, 215 with a FC ≤ 0.80 and 621 with a FC ≥ 1.20 (Fig. 1E). Functional annotation of the differentially abundant proteins, with KEGG pathway and gene ontology (GO) term enrichment analysis, identified multiple pathways of interest in *TAZ*^Δ45^ cells (Table S2A-B) (18, 19).

We identified 86 significantly enriched (p ≤ 0.05) KEGG pathways and GO terms for proteins with a FC ≤ 0.80, such as: oxidative phosphorylation (p=2.7 × 10^−6^), mitochondrial respiratory chain complex I assembly (p=7.9 × 10^−5^), mitochondrial chain complex I (p= 2.1 × 10^−4^), response to oxidative stress (p=1.6 × 10^−3^), NADH dehydrogenase (ubiquinone) activity (p=2.1 × 10^−3^), and metabolic pathways (p=0.019) (Table S2A). We identified 127 significantly enriched (p ≤ 0.05) KEGG pathways and GO terms for proteins with a FC ≥ 1.20, such as: mitochondrion (p=2.2 × 10^−4^), metabolic pathways (p=3.1 × 10^−3^), positive regulation of apoptotic process (p=4.2 × 10^−3^), AMPK signaling pathway (p=7.5 × 10^−3^), response to oxidative stress (p=9.3 × 10^−3^)(Table S2B).

### Functional annotation analysis: proteins of respiratory complex I and mitochondrial quality control

Defects in OXPHOS function have been previously described in BTHS, and consistent with these previous studies, we found that of the 86 terms significantly enriched for proteins with FC ≤ 0.80 in *TAZ*^Δ45^ cells, 18 reference the mitochondrion and/or OXPHOS, including the OXPHOS KEGG pathway (Table S2A). Specifically, 11 proteins with a FC ≤ 0.80 are encoded by genes associated with the OXPHOS KEGG pathway (Fig. S3). Of these 11 proteins, 5 are subunits of complex I (CI) and the remaining 6 proteins are subunits of complex III, IV, and V(Fig. S3). When we further examined the 18 terms that reference the mitochondrion and/or OXPHOS, we found that 4 specifically reference CI of OXPHOS (Table S2A).

The enrichment of the CI-associated terms in our functional annotation analysis was driven by 7 proteins encoded by the genes: *MT-ND3, NDUFAF1, NDUFA5, NDUFAB1, NDUFB2, NDUFB4,* and *OXA1L* (Fig. 2A). Five are subunits of CI (*MT-ND3, NDUFA5, NDUFAB1, NDUFB2, NDUFB4*), one is a CI assembly factor (*NDUFAF1*), and one assists with inserting proteins into the mitochondrial membrane and has been implicated in CI biogenesis (*OXA1L*) (20, 21). In total, proteomics identified and quantified 56 CI associated proteins in WT and *TAZ*^Δ45^ cells, and 45/56 had reduced abundance in *TAZ*^Δ45^ cells (FC range=0.608-0.998) (Table S3). Together, the functional annotation analysis highlights a decreased abundance of proteins associated with complex I, further delineating a previously described pathway in BTHS.

**Figure 2.**
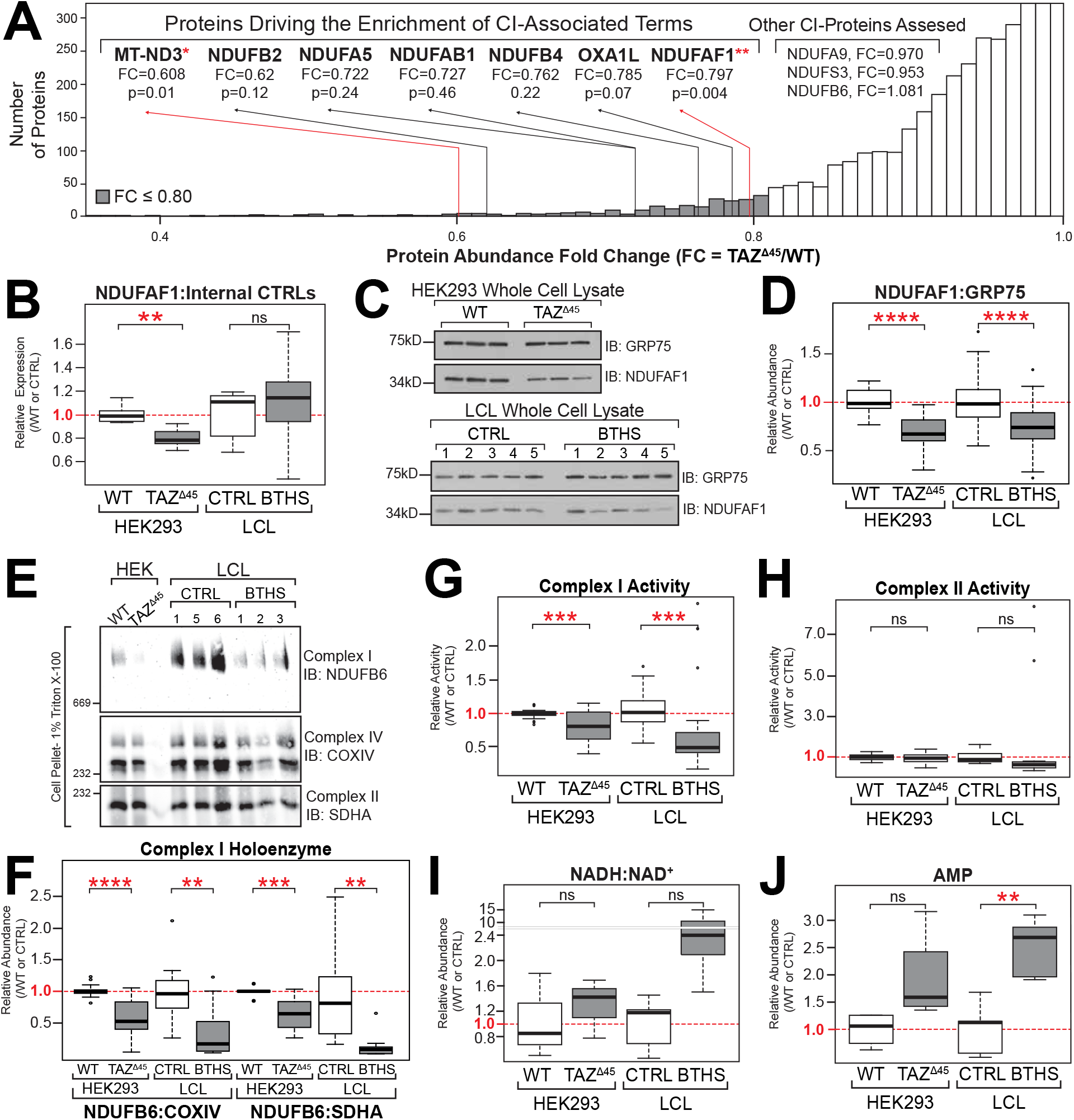
Reduced complex I (CI) holoenzyme abundance and activity in HEK293 TAZ^Δ45^ cells and BTHS patient-derived lymphoblastoid cells. **(A)** The enrichment of the CI-associated terms is driven by 7 genes; *MT-ND3, NDUFAF1, NDUFA5, NDUFAB1, NDUFB2, NDUFB4, OXA1L*. **(B)** Relative mRNA expression of *NUDFAF1*, determined by qRT-PCR and ΔΔCT quantification. Data plotted relative to WT/CTRL expression; WT n=6, *TAZ*^Δ45^ n=6, CTRL n=10, BTHS n=15. **(C)** Whole cell lysate (45 ug) of the indicated lines were immunoblotted for the indicated proteins. **(D)** Band intensities, relative to the loading control GRP75, were quantified and plotted relative to WT/CTRL abundance; WT n=27, *TAZ*^Δ45^ n=26, CTRL n=49, BTHS n=48. **(E)** BN-PAGE of HEK293 WT and *TAZ*^Δ45^ or CTRL and BTHS LCL cells (100-250K cells) solubilized in 1% Triton X-100 immunoblotted for the indicated proteins. **(F)** Band intensities were quantified, and CI abundance was represented as the ratio of CI band intensity to CIV (COXIV) or CII (SDHA). Abundance was plotted relative to WT/CTRL abundance; CI:CIV(WT n=13, *TAZ*^Δ45^ n=16, CTRL n=12, BTHS n=12), CI:CII (WT n=10, *TAZ*^Δ45^ n=13, CTRL n=9, BTHS n=9). **(G)** CI activity measured in HEK293 WT and *TAZ*^Δ45^ or CTRL and BTHS LCLs mitochondria (200 ug total protein) on a microplate reader (450nm) by following the oxidation of NADH to oxidized nicotinamide adenine dinucleotide (NAD+). Activity plotted relative to WT/CTRL abundance; WT n=25, *TAZ*^Δ45^ n=26, CTRL n=36, BTHS n=42. **(H)** CII activity measured in HEK293 WT and *TAZ*^Δ45^ or CTRL and BTHS LCLs mitochondria (200 ug total protein) on a microplate reader by following the production of ubiquinol by CII coupled to the reduction of the dye diclorophenolindophenol (DCPIP, 600nm). Activity plotted relative to WT/CTRL abundance; WT n=12, *TAZ*^Δ45^ n=12, CTRL n=10, BTHS n=12. Targeted metabolomics was used to measured **(I)** NADH, NAD+, and **(J)** cellular AMP via mass spectrometry in HEK293 WT (n=3), HEK293 *TAZ*^Δ45^ (n=3), CTRL LCL (n=5) and BTHS LCL (n=5) cells. Significant differences are indicated; * ≤ 0.05, ** ≤ 0.005, *** ≤ 0.0005, **** ≤ 0.00005.

When we analyzed the top five proteins with significantly increased abundance, we identified PARL as a candidate for further study due to its role in regulating mitochondrial responses to stress, such as membrane depolarization and increased reactive oxygen species (22–24). Additionally, dysregulation of PARL substrates has been implicated in cardiomyopathy and cardiac development, further highlighting the potential role of PARL dysregulation in BTHS (25–27). Upon further investigation of the 127 terms significantly enriched for proteins with a FC > 1.20, we observed that 20 terms reference the mitochondrion or mitochondrial dynamics, including: metabolic pathways (p=3.1 × 10^−3^), positive regulation of the apoptotic process (p=4.2 × 10^−3^), and mitochondrial inner membrane (p=5.2 × 10^−3^) (Table S2B). The functional annotation analysis revealed other genes of interest due to their potential roles in apoptosis, lipid trafficking, and/or mitochondrial quality control (MQC), such as; PRELID1, PRELID3B, CASP2, CASP7, CASP8, and CASP9 (28–30). Together with the proteomics findings, the functional annotation analysis suggests an increased abundance of proteins involved in MQC, a pathway not previously described in BTHS.

### Reduced complex I holoenzyme and activity in HEK293 *TAZ*^Δ45^ cells and BTHS patient derived lymphoblastoid cells

To assess the biological significance of reduced CI associated proteins identified via proteomics, we further investigated CI subunit/holoenzyme mRNA expression, protein abundance, and function. We measured the relative mRNA expression of *NDUFAF1*, the most significantly reduced protein identified by proteomics (FC=0.797, p=0.004), in both HEK293 and BTHS patient derived lymphoblastoid cells (LCLs). In *TAZ*^Δ45^ cells, *NDUFAF1* had reduced mRNA expression (0.80, p=6.4 × 10^−4^), whereas in BTHS LCLs there was no significant difference in mRNA expression (Fig. 2B).

We also measured the relative mRNA expression the 5 other CI associated proteins that had a FC ≤ 0.80 (MT-ND3, NDUFA5, NDUFB2, NDUFAB1, and NDUFB4), as well as 3 additional CI associated proteins (NDUFA9, NDUFS3, and NDUFB6), which did not have a FC≤ 0.80 but had been previously shown to have reduced protein abundance in BTHS LCLs (Fig. 2A & Fig. S4) (17). Of these 8 genes, 4 had significantly reduced mRNA expression in *TAZ*^Δ45^ cells (*NDUFB2* p=0.04, *NDUFAB1* p=0.001, *NDUFB4* p=0.02, and *NDUFB6* p=0.01) (Fig. S4).*NDUFA5, NDUFA9,* and *NDUFS3* had reduced mRNA expression in *TAZ*^Δ45^ cells but this reduction did not reach statistical significance (Fig. S4). There were no significant differences in the mRNA expression between control and BTHS LCLs, however there was significant variability in mRNA expression of all genes between the 5 individual BTHS LCL lines (Fig. S5). These individual differences could be due to the individual patient’s genetic background or the EBV transformation of the LCLs and may obfuscate the biological significance of mRNA expression in patient-derived LCLs.

Immunoblotting of HEK293 and LCL whole cell lysate for NDUFAF1 confirmed reduced protein abundance in *TAZ*^Δ45^ cells (FC=0.69, p=4.89 × 10^−10^) and BTHS LCLs (FC=0.75, p=7.69 × 10^−7^) (Fig. 2C). We also immunoblotted HEK293 and LCL mitochondria for the 3 additional CI subunits that had been previously shown to have reduced protein abundance (17). There was a subtle trend towards reduced NDUFA9, NDUFS3, and NDUFB6 abundance in *TAZ*^Δ45^ cells (p=ns, p=0.04, p=0.05, respectively), and a strong reduction in BTHS LCLs (p=7.6 × 10^−6^, p=1.8 × 10^−7^, p=5.1 × 10^−6^ respectively) (Fig. S6). The abundance of NDUFA9, NDUFS3, and NDUFB6 in *TAZ*^Δ45^ cells aligns with our proteomics findings, that found no significant difference in the abundance of these proteins between WT and *TAZ*^Δ45^ cells (Fig. 2A, Fig. S6). In order to determine whether TAZ deficiency affected the protein abundance of subunits from other respiratory complexes, we also immunoblotted for UQCRC2, a subunit of respiratory complex III. We found no significant difference in UQCRC2 abundance between WT and *TAZ*^Δ45^ cells, and a subtle but significantly reduced abundance of UQCRC2 in BTHS LCLs compared to controls (Fig. S6).

To determine the total abundance of CI holoenzyme, HEK293 and LCL cells were solubilized with Triton X-100 and resolved by BN-PAGE for the quantification of individual respiratory complexes. CI was the most significantly reduced complex, and the ratio of CI to CIV (CI:CIV) or CI:CII was significantly reduced in both *TAZ*^Δ45^ vs. WT (p=2.32 × 10^−5^, p=1.62 × 10^−4^, respectively) and BTHS LCLs vs. controls (p=0.001, p=0.005, respectively) (Fig. 2D). By comparing starting material and the cellular pellet following solubilization via immunoblotting, we found no significant difference in the solubilization of *TAZ*^Δ45^ and WT cells (87% and 88% solubilization efficiency, respectively). There was a minimal though statistically significant difference in the solubilization of BTHS and control LCLs (84% and 79% solubilization efficiency, respectively, p=0.03). Therefore, the observed differences were not due to an effect of altered CL on Triton X-100 solubilization.

Using a colorimetric CI enzyme activity assay that detects the oxidation of NADH to NAD^+^, we found a significant reduction in CI activity in both *TAZ*^Δ45^ cells and BTHS LCLs (p=9.4 × 10^−5^ & p=1.8 × 10^−4^, respectively) (Fig. 2E). Using a colorimetric CII enzyme activity assay, we found no significant difference in CII activity between WT and *TAZ*^Δ45^ cells or between control and BTHS LCLs, emphasizing the preeminent role of CI in BTHS-associated OXPHOS dysfunction (Fig. 2F). There was a wide range of CII function in LCLs derived from different individuals (Fig. S7). Overexpression of NDUFAF1, a CI assembly factor and the most significantly reduced CI associated protein in *TAZ*^Δ45^ cells, did not normalize the CI functional deficiency in *TAZ*^Δ45^ cells, indicating that CI dysfunction is due to a combination of reduced subunits and assembly factors (Fig. S8).

Measurement of intracellular NADH and NAD+ showed a trend towards an increase in the ratio of NADH to NAD^+^ in both *TAZ*^Δ45^ and BTHS LCLs compared to either WT or controls, though neither reached statistical significance (Fig. 2G). There was also an increase in intracellular AMP in both *TAZ*^Δ45^ cells and BTHS LCLs compared to either WT or controls, with a significant increase in the BTHS LCLs (p=0.002) (Fig. 2H). This moderate disturbance in energy homeostasis is consistent with the modest perturbations we observed in CI from mRNA expression, to protein expression, and ultimately activity in TAZ-deficient cells.

### Increased PARL abundance correlates with increased cleavage of downstream target PGAM5

To confirm the proteomics finding of increased PARL abundance (FC=1.815, p=0.016) in TAZ-deficient HEK293 cells, we immunoblotted HEK293 and LCL whole cell lysate for PARL. Using a PARL-KO HEK293 cell line generously provided by the Langer Laboratory, we identified a single band at ∼33kD present in both WT and *TAZ*^Δ45^ cells and absent in the PARL-KO cells, consistent with the band representing mature mitochondrial PARL (MAMP-PARL), referred to as PARL (Fig. S9) (24).

In *TAZ*^Δ45^ cells there was a significant increase in the abundance of PARL (FC=1.51, p=1.81 × 10^−10^) (Fig. 3A). There was a subtle but significant increase in the relative mRNA expression of *PARL* in *TAZ*^Δ45^ cells (WT=0.97, *TAZ*^Δ45^=1.09, p=0.02) (Fig. 3B). Immunoblotting of LCL whole cell lysate for PARL showed a significant reduction in PARL abundance, however, there was extreme variability in PARL abundance in the CTRL lines (CTRL 1-3 vs CTRL 4-5) (Fig. S10A). There was no significant difference in the mRNA expression of PARL between CTRL and BTHS LCLs or across the 5 different BTHS LCL lines (Fig. S10B-C). Thus, increased PARL in TAZ-deficient HEK293 cells may be regulated both at the transcriptional and protein expression levels, and lack of consistent findings in BTHS LCLs may be due to cell-type specific differences in PARL regulation or difficulty with detection in a small sample size due to high intra-subject variability.

**Figure 3.**
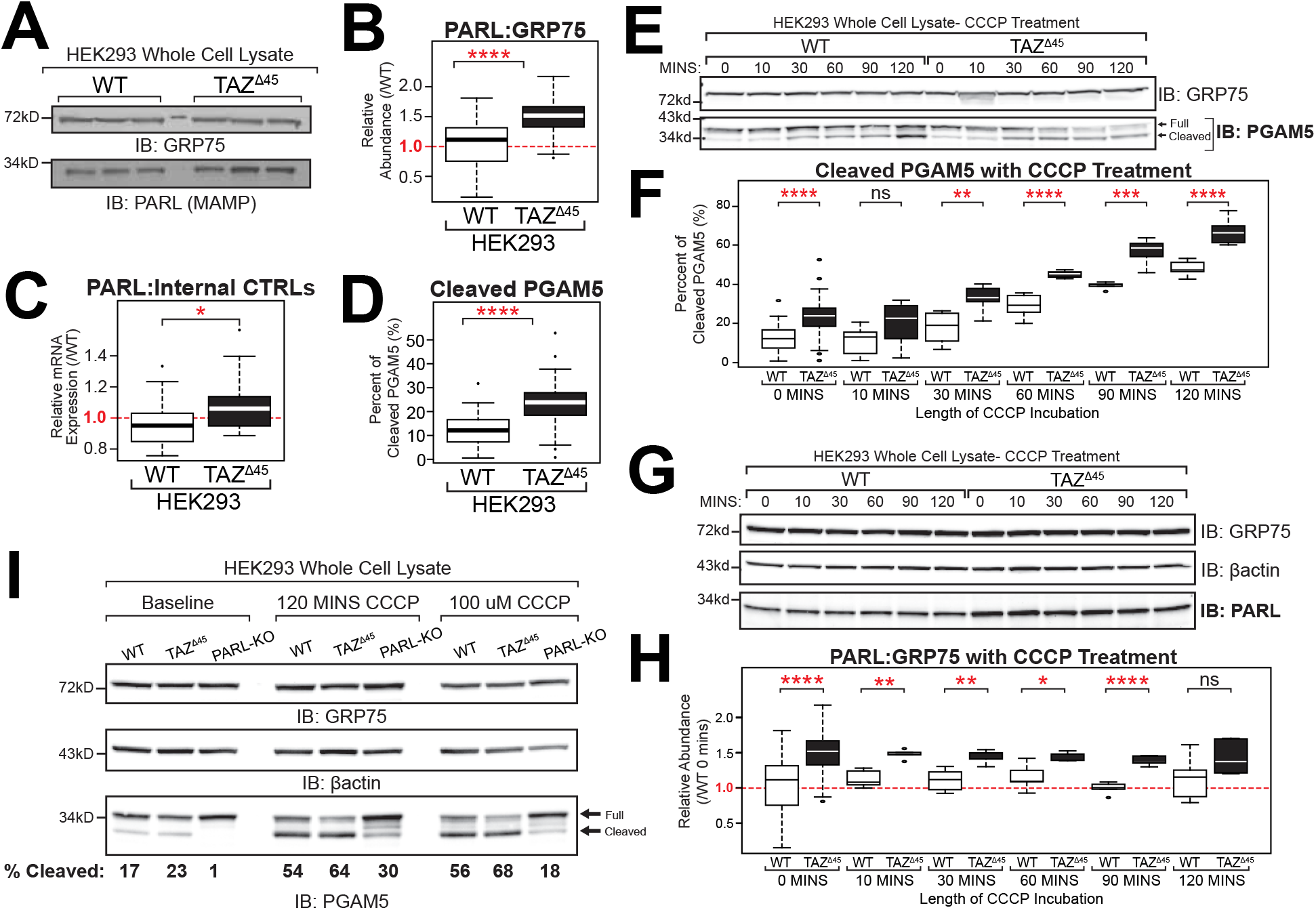
Increased cleavage of PGAM5 by PARL in HEK293TAZ^Δ45^ cells. **(A)** Whole cell lysate (45 ug) of the indicated lines were immunoblotted for the indicated proteins. **(B)** Band intensities, relative to loading control GRP75, were quantified and plotted relative to WT abundance; WT n=54, *TAZ*^Δ45^ n=48 **(C)** Relative mRNA expression of *PARL* determined by qRT-PCR and ΔΔCT quantification; WT n=18, *TAZ*^Δ45^ n=17. **(D)** Whole cell lysate (45 ug) of WT and *TAZ*^Δ45^ cells were immunoblotted for PGAM5 and loading control GRP75. Band intensities, relative to the loading control GRP75, for both full-length and cleaved PGAM5 were individually quantified and plotted as the percent of cleaved PGAM5 (cleaved/full+cleaved); WT n=41, *TAZ*^Δ45^ n=41. **(E)** HEK293 WT and *TAZ*^Δ45^ cells were treated with 20uM CCCP for serial time points from 0 to 120 minutes. Whole cell lysate (45 ug) of the indicated lines and treatment times were immunoblotted for the indicated proteins. **(F)** Band intensities, relative to the loading control GRP75, for both full-length and cleaved PGAM5 were individually quantified and plotted as the percent of cleaved PGAM5 (cleaved/full+cleaved); WT 0 mins n=41, WT 10 mins n=8, WT 30 mins n=8, WT 60 mins n=8, WT 90 mins n=5, WT 120 mins n=7, *TAZ*^Δ45^ 0 mins n=41, *TAZ*^Δ45^ 10 mins n=8, *TAZ*^Δ45^ 30 mins n=7, *TAZ*^Δ45^ 60 mins n=5, *TAZ*^Δ45^ 90 mins n=7, *TAZ*^Δ45^ 120 mins n=7. **(G)** HEK293 WT and *TAZ*^Δ45^ cells were treated with 20uM CCCP for serial time points from 0 to120 minutes. Whole cell lysate (45 ug) of the indicated lines and treatment times were immunoblotted for the indicated proteins. **(H)** Band intensities, relative to the loading control GRP75, were quantified and plotted relative to WT abundance; WT 0 mins n=54, WT 10 mins n=5, WT 30 mins n=6, WT 60 mins n=6, WT 90 mins n=5, WT 120 mins n=6, *TAZ*^Δ45^ 0 mins n=48, *TAZ*^Δ45^ 10 mins n=5, *TAZ*^Δ45^ 30 mins n=5, *TAZ*^Δ45^ 60 mins n=5, *TAZ*^Δ45^ 90 mins n=5, *TAZ*^Δ45^ 120 mins n=6. **(I)** Whole cell lysate (45 ug) of the indicated lines were immunoblotted for the indicated proteins. Band intensities, relative to the loading control GRP75, for both full-length and cleaved PGAM5 were individually quantified, and the percent of cleaved PGAM5 (cleaved/full+cleaved) is indicated for each lane (n=1). Significant differences are indicated; * ≤ 0.05, ** ≤ 0.005, *** ≤ 0.0005, **** ≤ 0.00005.

To investigate the biological significance of increased PARL abundance in *TAZ*^Δ45^ cells, we assessed a downstream proteolytic target of PARL, PGAM5. Previous evidence suggests that PGAM5 is cleaved by PARL and another stress-activated IMM protease, OMA1, in response to loss of mitochondrial membrane potential (ΔΨm) (31). At baseline, we observed a significant increase in the percent of cleaved PGAM5 in *TAZ*^Δ45^ cells (p=1.5 × 10^−7^) (Fig. 3C). There was no significant difference in the percent of cleaved PGAM5 between CTRL and BTHS LCLs (Fig. S10D)

Upon treatment with membrane depolarizer carbonyl cyanide m-chlorophenyl hydrazine (CCCP), with serial time point measurements, we observed that *TAZ*^Δ45^ cells maintained a significantly greater abundance of PARL until the final timepoint of 120 mins (Fig. 3E, Table S5). This increase in PARL abundance with CCCP correlates with the increase in PGAM5 cleavage observed in both WT and *TAZ*^Δ45^ cells (Fig. 3D). Under the same CCCP treatment protocol, we also observed an increase in the percentage of cleaved PGAM5 in WT cells and *TAZ*^Δ45^ cells, where *TAZ*^Δ45^ cells maintained a significantly greater percentage of cleaved PGAM5 at all time points tested (Fig. 3D). The difference in the percentage of cleaved PGAM5 between WT and *TAZ*^Δ45^ cells increased over time, from a 11% difference at 0 mins to a 18% difference at 120 mins (Fig. 3D, Table S4).

We further demonstrated that PGAM5 cleavage was absent and reduced inn PARL-KO cells not treated or treated with CCCP, respectively, underscoring that PGAM5 is predominantly cleaved by PARL (Fig. 3F). In summary, we observed baseline abnormalities in PARL abundance and PGAM5 cleavage, which is exacerbated upon mitochondrial depolarization, in TAZ-knockout HEK293 cells.

### Targeting CL with bromoenol lactone and SS-31 normalizes downstream cellular dysfunction

To determine if targeting CL and CL metabolism would affect the dysregulation observed in respiratory CI and/or MQC, *TAZ*^Δ45^ and WT cells were treated with either bromoenol lactone (BEL) or SS-31. Previous studies have shown that treatment with BEL, an inhibitor of calcium-independent PLA2 (iPLA2), partially remediates CL abnormalities by reducing MLCL accumulation and CL depletion (32, 33). SS-31, a cell permeable mitochondria-targeted tetrapeptide, selectively binds CL where it has been shown to stabilize cristae morphology and preserve mitochondrial bioenergetics (34, 35).

The relative abundance of NDUFAF1 in *TAZ*^Δ45^ cells increased following BEL treatment and significantly increased following SS-31 treatment (p= 2.8 × 10^−5^) (Fig. 4A, Table S6). *NDUFAF1* relative mRNA expression significantly increased in *TAZ*^Δ45^ cells following both BEL and SS-31 treatment (p= 0.05 and p= 0.01, respectively) (Fig. 4B, Table S7). The relative mRNA expression was also measured for the other 4 CI associated genes that had significantly reduced levels in *TAZ*^Δ45^ cells at baseline (*NDUFB2, NDUFAB1, NDUFB4*, and *NDUFB6*). Following BEL treatment, these significant differences in expression between *TAZ*^Δ45^ vs. WT were no longer observed in any of the 4 genes tested, and following SS-31 treatment, the significant differences were no longer observed in 3 of the 4 genes tested (Fig. S12, Table S7).

**Figure 4.**
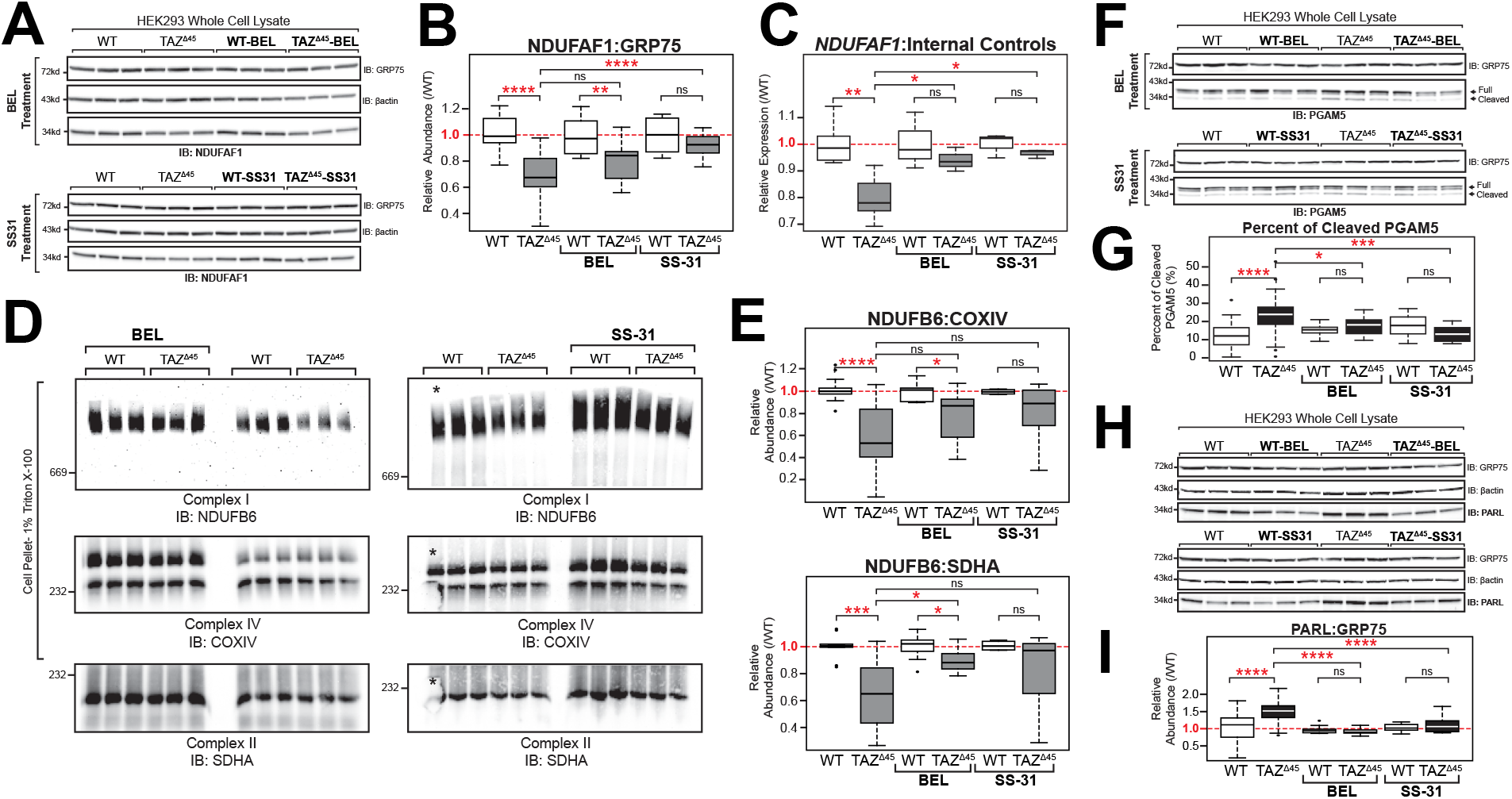
Targeting CL or CL metabolism with bromoenol lactone and SS-31 modifies mitochondrial dysfunction. **(A)** HEK293 WT and *TAZ*^Δ45^ cells were treated for 48 hours with 2.5uM bromoenol lactone (BEL) and 7 days with 100nM SS-31. Whole cell lysate (40-45 ug) of the indicated lines were immunoblotted for the indicated proteins. **(B)** Band intensities, relative to the loading control GRP75, were quantified and plotted relative to WT abundance; WT n=27, *TAZ*^Δ45^ n=26, WT-BEL n=9, *TAZ*^Δ45^-BEL n=9, WT-SS-31 n=9, *TAZ*^Δ45^-SS-31 n=9. **(C)** Relative mRNA expression of *NDUFAF1* determined by qRT-PCR and ΔΔCT quantification using each respective control; WT n=6, *TAZ*^Δ45^ n=3, WT-BEL n=3, *TAZ*^Δ45^-BEL n=3, WT-SS-31 n=3, *TAZ*^Δ45^-SS-31 n=3 per gene. **(D)** BN-PAGE of HEK293 WT and *TAZ*^Δ45^ cells treated for 48 hrs with 2.5uM BEL (120 ug total protein) and 7 days with 100nM SS-31 solubilized in 1% Triton X-100 immunoblotted for the indicated proteins. All samples were resolved on a single gel and exposed for the same duration. The WT lane indicated with the asterisk was not used for quantification due to air bubbles. **(E)** Band intensities were quantified, and CI abundance was represented as the ratio of CI band intensity to CIV or CII. Abundance was plotted relative to respective control; CI:CIV(WT n=13, *TAZ*^Δ45^ n=16, WT-BEL n=9, *TAZ*^Δ45^-BEL n=11, WT-SS-31 n=4, *TAZ*^Δ45^-SS-31 n=6), CI:CII (WT n=10, *TAZ*^Δ45^ n=13, WT-BEL n=9, *TAZ*^Δ45^-BEL n=10, WT-SS-31 n=4, *TAZ*^Δ45^-SS-31 n=6). **(F)** Whole cell lysate (40-45 ug) of the indicated lines and treatment conditions were immunoblotted for the indicated proteins. **(G)** Band intensities, relative to the loading control GRP75, for both full-length and cleaved PGAM5 were individually quantified and plotted as the percent of cleaved PGAM5 (cleaved/full+cleaved); WT n=41, *TAZ*^Δ45^ n=41, WT-BEL n=16, *TAZ*^Δ45^-BEL n=16, WT-SS-31 n=9, *TAZ*^Δ45^-SS-31 n=8. **(H)** Whole cell lysate (40-45 ug) of the indicated lines and treatment conditions were immunoblotted for the indicated proteins. **(I)** Band intensities, relative to the loading control GRP75, were quantified and plotted relative to WT abundance; WT n=54, *TAZ*^Δ45^ n=48, WT-BEL n=11, *TAZ*^Δ45^-BEL n=11, WT-SS-31 n=10, *TAZ*^Δ45^-SS-31 n=12. Significant differences are indicated; * ≤ 0.05, ** ≤ 0.005, *** ≤ 0.0005, **** ≤ 0.00005.

Next, CI holoenzyme abundance was measured in BEL and SS-31 treated cells by BN-PAGE. CI remained the most significantly reduced complex in *TAZ*^Δ45^ cells after BEL or SS-31 treatment (Fig. 4C). However, both treatments had a subtle effect on the ratio of both CI to CIV (CI:CIV) and CI to CII (CI:CII) (Fig. 4C, Table S8). The CI:CIV relative abundance increased from 0.59 in untreated *TAZ*^Δ45^ cells to 0.77 in *TAZ*^Δ45^-BEL cells and 0.80 in *TAZ*^Δ45^-SS-31 cells (Fig. 4C, Table S8). The CI:CII relative abundance significantly increased from 0.66 in untreated *TAZ*^Δ45^ cells to 0.89 in *TAZ*^Δ45^-BEL cells (p= 6.1 × 10^−3^) and to 0.83 in *TAZ*^Δ45^-SS-31 cells (Fig.4C, Table S8). Overall, treatment with either BEL or SS-31 showed a subtle increase in CI holoenzyme abundance.

Immunoblotting of BEL and SS-31 treated HEK293 whole cell lysate for PGAM5 showed a significant decrease in the percentage of cleaved PGAM5 in *TAZ*^Δ45^-BEL (18%, p= 0.01) and *TAZ*^Δ45^-SS-31 (13%, p= 2.6 × 10^−4^) cells compared to TAZ^Δ45^-untreated cells (23%) (Fig. 4D, Table S9). There was no significant difference in the percentage of cleaved PGAM5 in *TAZ*^Δ45^-BEL versus WT-BEL cells or *TAZ*^Δ45^-SS-31 versus WT-SS-31 cells (Fig. 4D, Table S9). Further, immunoblotting of BEL and SS-31 treated HEK293 whole cell lysate for PARL showed a significant decrease in the relative abundance of PARL in *TAZ*^Δ45^-BEL (FC= 0.93, p= 7.1 × 10^−15^) and *TAZ*^Δ45^-SS-31 (FC= 1.11, p= 9.9 × 10^−5^) cells compared to TAZ^Δ45^-untreated cells (FC= 1.53) (Fig. 4E, Table S10), which was essentially restored to WT levels. Collectively, these results indicate that drugs that target CL partially rescue the defects in CI and MQC observed in TAZ-deficient cells.

## Discussion

As a central phospholipid of the IMM, CL has been shown to have diverse roles in mitochondrial function (1). Yet, these diverse roles are currently underappreciated in the pathophysiology of Barth syndrome, which is the only known Mendelian disorder of CL metabolism. In this study, we took an untargeted approach to identify dysregulated proteins in TAZ-deficient HEK293 cells and pursued two areas of dysregulation for further study: CI of the respiratory chain and components of mitochondrial quality control.

Previous work has linked TAZ deficiency and mitochondrial respiratory chain dysfunction, with several studies pointing towards CI as the most impacted respiratory chain complex (15, 17, 36). In this work we confirmed that CI expression and function are abnormal in TAZ deficiency. We further showed that this dysfunction is potentially driven by the decreased expression of 5 subunits of mitochondrial CI, *MT-ND3, NDUFA5, NDUFAB1, NDUFB2, NDUFB4,* and reduced expression of the mitochondrial assembly factor, *NDUFAF1*. These specific deficiencies could disrupt aspects of the coordinated modular assembly of CI and further studies into CI modular assembly in BTHS could prove to be revealing (20, 37).

We further identified abnormal expression and regulation of MQC associated proteins including PARL and PGAM5. MQC, involving the processes of mitochondrial proteostasis, biogenesis, dynamics, and mitophagy, is emerging as a central theme for the pathogenesis of various diseases (38–40). PARL participates in MQC via reciprocal proteolysis of PGAM5 and PINK1 (22, 31). Upon mitochondrial depolarization, PARL upregulation accelerates PGAM5 proteolysis which drives mitochondrial fragmentation (31). Defective mitochondrial quality control, particularly as it affects cellular energy production and stress responsiveness, is increasingly recognized for its role in diverse forms of cardiac dysfunction and may be central to the cardiac pathogenesis of BTHS (41).

In order to determine if the abnormalities we identified are directly associated with abnormal CL, we tested the ability of a pharmacological inhibitor of PLA2γ (BEL), a protein capable of generating MLCL from CL, and a CL binding peptide (SS-31) to ameliorate the two TAZ-deficient phenotypes we established. Indeed, we showed that targeting CL and CL metabolism with BEL and SS-31 is sufficient to modulate CI subunit gene expression and protein abundance, PARL abundance, and PGAM5 cleavage. Thus, targeting multiple aspects of CL metabolism may be a feasible therapeutic approach in BTHS.

In terms of future directions for study, it is notable that among the CI subunits that we identified as having altered expression in TAZ deficiency, NDUFAB1 has an essential role in fatty acid biosynthesis due to its dual role as the mitochondrial acyl carrier protein (mt-ACP) (42). In fact, the GO term “protein lipoylation” was identified in the functional annotation of proteins with a FC ≤ 0.80 (Table S2A). In this role, abnormal expression of NDUFAB1/mt-ACP in TAZ deficiency may have further implications beyond affecting CI assembly and function, including affecting CL acyl chain content. Additionally, mt-ACP is involved in the assembly of the mitochondrial ribosome, and therefore mt-ACP expression and abundance may further influence changes in mitochondrial gene expression (43). Further examining the essential role of NDUFAB1, outside of CI assembly and function, may provide a mechanistic link between altered CL metabolism and altered bioenergetic metabolism. Defects in MQC pathways, have been implicated in the pathogenesis of various cardiac pathologies relevant to BTHS. Additionally, enlarged mitochondria have been observed in BTHS patient and mouse model derived cardiac tissue, consistent with impaired mitophagy (44–47). Thus, further examining these pathways in an appropriate cellular context could provide insight into potential therapeutic targets for BTHS and other conditions resulting from MQC dysfunction.

Finally, it is important to note that some of our findings were cell-type specific (HEK293 vs. LCL). We found that HEK293s and LCLs differed in PARL abundance, which is consistent with previous studies confirming cell-type specific expression and regulation of PARL, and introduces further questions about whether cell-type specific mechanisms of MQC contribute to the tissue distribution of disease in BTHS (48). We also identified a reduction in CI subunit protein abundance, holoenzyme abundance, and CI activity in both HEK293 TAZ-KO cells and BTHS LCLs; however, in BTHS LCLs reduced mRNA expression of CI subunits was not observed. The differences in mRNA expression observed between the HEK293s and the LCLs could be due to epigenetic and transcriptomic differences observed in EBV transformed LCL lines, the genetic differences between the BTHS individuals, and/or cell type specific regulation of CI associated genes/proteins (49). This hypothesis emphasizes the need for a clearer understanding of the cell-specific effects of abnormal CL content. To explore these questions, we are presently exploring respiratory chain and MQC dysfunction in diverse TAZ-KO cell types in order to understand how these pathways are associated with the tissue specific expression of BTHS.

## Materials and Methods

### Cell lines and culture conditions

HEK293 WT cells were purchased from ATCC (293 [HEK-293]x ATCC® CRL-1573(tm)). Collection of control (LCL Control #1) and BTHS patient (LCL BTHS #1-5) derived LCL lines had institutional IRB approval via Johns Hopkins University protocols #IRB00098987 (Table S11). Individuals were diagnosed with BTHS via an increased MLCL:CL ratio. Additional control LCL lines (n=9) were acquired through the following sources: Biochemical Genetics Laboratory at Kennedy Krieger Institute (LCL Control #2-3), Valle Laboratory (LCL Control #4-5), Coriell NINDS Biobank (LCL Control #6-9) (Table S11).

LCL lines were transformed by the Biochemical Genetics Laboratory at Kennedy Krieger Institute and the Valle Laboratory used the following protocol. Peripheral blood samples were centrifuged for 15 minutes at 3000 rpm. The “buffy coat” was then resuspended in RPMI and further centrifuged for 10 minutes at 1000 rpm. The resulting cell pellet was resuspended in RPMI and incubated with Epstein-Barr (EB) virus and T-cell growth factor (TCGF) at 37C for 48-72 hours. After incubation period, additional RPMI was added to the flask. Cells were monitored and fed RMPI for two weeks, after which the established transformed cell lines were passaged for experiments and/or freezing. LCL lines acquired through the Coriell NINDS Biobank were also transformed via EB virus.

All cells were grown at 37°C, 5% CO2. HEK293 WT and TAZ^Δ45^ were maintained in DMEM with L-glutamine and 4.5 g/L Glucose (Corning Cellgro Cat. #10-017) containing 10% fetal bovine serum (FBS, Gemini) and 2 mM L-glutamine (Gibco, Cat. #25030149). Patient derived lymphoblastoid cells (LCLs) were maintained in RPMI 1640 (Gibco, Cat. # 11875119) containing 10% FBS (Gemini). Mycoplasma contamination was routinely monitored and not detected.

For CCCP treatment, HEK293s were seeded into 6-well plates. At confluence, cells were either treated with 20uM CCCP (Cat. No) for 0,10,30,60,90, and 120 mins, or 0,10,20,40,80,100uM CCCP for 45 mins.

For BEL treatment, HEK293s (400K) were seeded into 6-well plates. 48hrs later, at 80-90% confluence, cells were treated with 2.5uM BEL (Cat. No) for 48hrs.

For SS-31 treatment, HEK293s (50K) were seeded into 6-well plates. For 7 days, cells were fed fresh 100nM SS-31.

### CRISPR/Cas9 genome editing

sgRNAs (Figure S1 & Table SI) targeting exon 2 of *TAZ* were designed at www.crispr.mit.edu and selected based on the scoring algorithm detail in Hsu et al. 2013 (Table S1) (50, 51). Synthesized sgRNA 1 and sgRNA 2 were individually cloned into pSpCas9(BB)-2A-Puro (PX459) V2.0 vector pSpCas9(BB)-2A-Puro (PX459) V2.0 was a gift from Feng Zhang (Addgene plasmid # 62988; http://n2t.net/addgene:62988; RRID:Addgene_62988 (50). HEK293 WT cells were transfected with both sgRNA vectors using Lipofectamine 2000 (Invitrogen). 24-hours after transfection cells were subjected to Puromycin selection (2ug/mL) for approximately 48 hours. Following selection, cells we passaged in order to isolate single cell clones. Confluent colonies of single cell clones were collected using a cloning cylinder and expanded for DNA isolation and screening.

### Screening and genotyping

Genomic DNA was extracted from an aliquot of 3 × 10^6^ cells using the Gentra Purgene Kit (Qiagen). The genomic region surrounding the CRISPR/Cas9 target site (741 bp) was PCR amplified using AccuPrime GC-Rich DNA Polymerase (Invitrogen) (Table S1). The PCR product was then used for screening with Surveyor Assay Kit (IDT). The PCR product of Surveyor-Positive clones was further analyzed by Sanger sequencing.

### RT-PCR

RNA was extracted from an aliquot of 3 × 10^6^ cells using the RNeasy Plus Kit (Qiagen). cDNA was synthesized from extracted RNA using the iScript cDNA Synthesis Kit (BioRad). The region surrounding the CRISPR/Cas9 target site was PCR amplified using AccuPrime GC-Rich DNA Polymerase (Invitrogen) (Forward Primer: 5’ TACATGAACCACCTGACCGT 3’, Reverse Primer: 5’ CAGATGTGGCGGAGTTTCAG 3’). PCR products were resolved on a 0.8% agarose gel.

### Whole Cell Lysate Extraction

An aliquot of 3 × 10^6^ cells was centrifuged for 4 minutes at 1000rpm. The resultant cell pellet washed twice with ice-cold PBS and lysed with RIPA lysis buffer (1% (v/v) Triton X-100, 20 mm HEPES–KOH, pH 7.4, 50 mm NaCl, 1 mm EDTA, 2.5 mm MgCl2, 0.1% (w/v) SDS) spiked with 1 mm PMSF for 30 min at 4°C with rotation. Insoluble material was removed by centrifugation for 30 min at 21000g at 4°C, the supernatant collected, and protein quantified using a bichichronic acid (BCA) assay (Pierce).

### Mitochondrial Isolation

Mitochondrial isolation of HEK293s was performed as previously described by Lu et al.(3). Mitochondrial isolation of LCLs was also performed as previously described, with slight modification. LCLs were grown in maintenance media until total cell count exceeded 100 × 10^6^ cells. LCLs were then centrifuged for 4 minutes at 1000rpm before beginning the previously described protocol by Lu et al.(3).

### Immunoblotting

Whole cell extracts or mitochondria, resuspended in XT Sample Buffer (BioRad) and Reducing Agent (BioRad), were resolved on Criterion XT 12% Bis-Tris gels (BioRad) in XT MOPS Running buffer (BioRad). Proteins were transferred to Immuno-Blot PVDF (BioRad) at 100V for 1 hour. Following transfer, membranes were blocked with 5% milk, 0.05% Tween-20 in PBS (Blocking Buffer) for 1 hour at room temperature or at 4C if longer, with rocking. Following blocking, membranes were briefly washed with PBST (PBS with 0.2% Tween-20) and then incubated with primary antibody in PBST with 0.02% Na-Azide overnight at 4C with rocking. Following three successive 10-minute washes with PBST at room temperature, HRP-conjugated secondary antibodies were added and incubated for 45 min at room temperature. The membranes were then washed three times for 10 minutes with PBST and twice for 10 minutes with PBS. Immunoreactive bands were visualized using ECL Western Blotting Detection (Amersham) or SuperSignal West Pico PLUS (Pierce). Images were captured with a Fluorchem Q (Cell Biosciences, Inc.) or on film. Film was scanned before quantification. Quantitation of bands was performed using ImageJ and protein expression values were normalized to loading controls.

### Antibodies

Antibodies against the following proteins were used; β-actin (loading control, Life Technologies #AM4302), GRP75 (loading control, 75-127), TAZ (#2C2C9)(3), NDUFAF1 (Abcam, #ab79826),PARL (Abcam #ab45231, Proteintech #26679-1-AP, Langer Laboratory), UQCRC2 (Abcam, #ab14745), NDUFA9 (Abcam, #ab14713), NDUFS3 (Abcam, #ab110246), NDUFB6 (Abcam,#ab110244). Two HRP-conjugated secondary antibodies were used; goat anti-rabbit (Abcam, #ab6721), goat anti-mouse (Abcam, #ab205719).

### Lipidomics

Lipids were extracted from cell pellets (3 × 10^6^ cells) and analyzed as previously described by Vaz et al. (52).

### Proteomics

Samples (HEK293 WT n=3, and TAZ^Δ45^ n=3, serial passages) were reduced in 5 mM DTT for 1 hour at 56C, alkylated in 10 mM iodoacetamide for 30 minutes in the dark at room temperature, and precipitated in cold (−20C) acetone 10% trichloroacetic acid. The precipitate was pelleted by centrifugation at 16,000 g for 15 minutes, the supernatant was discarded, and the pellet was rinsed with cold acetone and dried at room temperature. The samples were reconstituted in 50 µL of 20% acetonitrile 80 mM triethyl ammonium bicarbonate (TEAB) and digested with 3.3 µg of trypsin/Lys-C mix (Promega) at 37C overnight. The digested samples were labeled with TMT 10-plex reagent (Thermo, Lot #SK257889) and combined. The sample was lyophilized and reconstituted in 0.5 mL of 10 mM TEAB and fractionated by high (8-9) pH reversed phase chromatography into 84 fractions which were concatenated into 24. Each of the 24 fractions was lyophilized and reconstituted in 2% acetonitrile 0.1% formic acid and separated over a 90-minute low (2-3) pH reversed phase gradient (120 minutes run time) for mass spectrometry (MS) analysis on an Orbitrap Fusion.

MS data were acquired using serial data-dependent fragmentation of individual precursor ions (DDA). An intact precursor ion scan (MS1) spanning 400 - 1600 Th was acquired every 3 seconds at a resolution of 120,000 (at m/z = 200). Fragmentation scans (MS2) were acquired at 60,000 resolution between precursor scans by isolation of individual precursor ions at 0.6 Th resolution, accumulation to 5 x10^4^ automatic gain control for a maximum injection time of 250 ms, and activation with beam collision (HCD) at 38% energy. Mass accuracy was maintained throughout the experiment by internal calibrant lock mass.

The acquired data were searched against the SwissProt Human database by MASCOT using 4 ppm and 0.01 Da precursor and fragment maximum mass error, respectively. TMT labeled lysine and peptide N-termini and carbamidomethylation of cysteine were set as static modifications. Oxidation of methionine and deamidation of asparagine and glutamine were set as dynamic modifications. The results were rescored by Percolator in Proteome Discoverer 2.2 and quantitative analysis was carried out based on reporter ion intensities.

### Function Annotation

We selected proteins with a protein abundance fold change (FC, *TAZ*^Δ45^/ WT) less than or equal to 0.80 (FC ≤ 0.80) and proteins with a FC greater than or equal to 1.2 (FC ≥ 1.20). Each subset was individually uploaded to DAVID 6.8 and compared to the background “Homo sapiens” (18, 19). With the functional annotation tool, we pulled all KEGG pathways and GO terms for further analysis.

### Quantitative RT-PCR

Total RNA was extracted from an aliquot of 3 x 106 cells using the RNeasy Plus Kit (Qiagen). cDNA was synthesized from extracted RNA using the iScript cDNA Synthesis Kit (BioRad) in 20 μl reactions according to the manufacturer’s suggested protocol using 1 ug of RNA. cDNA was subsequently diluted 10-fold with water. 2.4 μl of cDNA was analyzed in 12 μl reactions using the SsoAdvanced Universal SYBR Green Supermix (Bio-Rad) according to the manufacturer’s instructions and included each respective forward and reverse gene-specific primers (Table S12). Each sample-primer reaction was plated in triplicate per plate. Each plate also included both no reverse-transcriptase controls (No-RT) for each cDNA sample and no template controls (No-Template) for each primer pair. The reaction conditions were as follows: 2 min at 95°C, followed by 40 two-temperature cycles of 5 s at 95°C and 30 s at 60°C. For nuclear genes, expression of each gene was analyzed by the comparative CT (ΔΔCT) method with *TBP* and *HPRT1* being averaged as endogenous reference genes. For mitochondrial genes (*MT-ND3)*, expression of each gene was analyzed by the comparative CT (ΔΔCT) method with *MT-RNR1, MT-CO1*, and *MT-ATP6* being averaged as endogenous reference genes. Values were represented as average fold change relative to respective wildtype or untreated controls.

### 1D BN-PAGE

Cell pellets (100,000 cells, ∼120ug) were solubilized for 30 min on ice in 20 mm Bis-Tris, 10% glycerol, 50 mM NaCl, supplemented with 1% (v/v) Triton X-100 and protease inhibitors (PMSF, Leupeptin, Pepstatin). Extracts were clarified by centrifugation for 30 min at 21 000g at 4C and analyzed by 1D BN/SDS–PAGE as described by Claypool et al. (53).

### Complex I and II activity assays

The activity of complex I was measured using the Complex I Enzyme Activity Microplate Assay Kit (Abcam, #ab109721) according to the manufacturer’s instructions using 200ug of isolated mitochondria for both HEK293s and LCLs. The activity of complex II was measured using the Complex II Enzyme Activity Microplate Assay Kit (Abcam, #ab109908) according to the manufacturer’s instructions using 200ug of isolated mitochondria for both HEK293s and LCLs.

### NDUFAF1 Transfection

HEK293 cells were seeded into 15 cm plates. When cells reached ∼80% confluency cells were transiently transfected with *NDUFAF1*(NM_016013) C-Myc/DDK-tagged plasmid (Origene #RC200029) with Lipofectamine 3000 (Thermo #L3000001) according to manufacturer’s instructions. 75.4 ug of plasmid was transfected with 105 uL of Lipofectamine 3000 per 15 cm plate. The cells were grown in galactose media for 48hrs before mitochondria extraction performed as described previously.

### Metabolomics

HEK and LCL cell samples undergoing metabolic analysis were initially kept on ice and washed with ice-cold PBS prior to collection, followed by centrifugation at 1500RPM and 4°C. Metabolites within the cell pellet were extracted with 80% HPLC-grade methanol (Fisher Scientific) and 20% mass-spec (MS)-grade water as previously described (54). The extraction solution was then collected and evaporated using a speed-vac and a lyophilizer which resulted in a white powder of dried metabolites. The collected metabolites were subsequently resuspended in 50% (vol/vol) acetonitrile diluted with MS-grade water and analyzed via an Agilent 1260 HPLC and 6490 triple-quadrupole (QQQ) mass spectrometer.

The Agilent 1260 HPLC-autosampler system was kept at 4°C for the entirety of the run to prevent any degradation within the samples. Optimal separation was achieved with a reverse phase chromatography system with an aqueous mobile phase of MS-grade water with 0.1% formic acid and an organic mobile phase of 98% acetonitrile with 0.1% formic acid. The Discovery® HS F5 HPLC Column (3µm particle size, L × I.D. 15 cm × 2.1 mm, Sigma) with a compatible guard column (Sigma) were used and maintained at a temperature of 35°C. The injection volume was 2µL and the runtime was 50 minutes per sample. The flow rate, buffer gradient, and mass spectrometer parameters for the method were the same as previously described (55).

Data from pure standards of each compound of interest were acquired prior and simultaneously with samples in identical setting: precursor and product ion transitions, collision energy, and ion polarity. Agilent MassHunter and Agilent Qualitative and Quantitative Analysis Software packages were used to analyze the metabolic profiles. The metabolite peaks were integrated for raw intensities and then normalized by protein concentration collected from the original cell pellet. Protein concentration was determined using a FilterMax F5 microplate reader and a Bovine Serum Albumin (BSA) standard.

### Data Analysis

All statistical analyses were performed using R version 3.3.2 (2016-10-31) (56). Between-group comparisons were performed using Welch’s two-sample t-test. Outliers, outside 1.5x the interquartile range above the upper quartile and below the lower quartile were only removed for statistical analyses of the isogenic HEK293 cell lines.

## Acknowledgements

We thank Ya-Wen Lu and Michelle Acoba of the Claypool Lab for their critical and fruitful technical guidance. We also want to thank Thomas Langer, and the Langer Laboratory for the generous gift of PARL antibody and PARL-KO HEK293 cells. Research reported in this publication was supported by the National Heart, Lung, And Blood Institute of the National Institutes of Health under Award Number F31HL147454 to A.F.A. The content is solely the responsibility of the authors and does not necessarily represent the official views of the National Institutes of Health. Proteomics analysis was supported by the Johns Hopkins University School of Medicine Core Coins Program to V.J.H.

