## Supplemental data for "Barth syndrome cellular models have dysregulated respiratory chain complex I and mitochondrial quality control due to abnormal cardiolipin"

**This PDF file includes:**

Figures S1 to S12

Tables S1 to S12

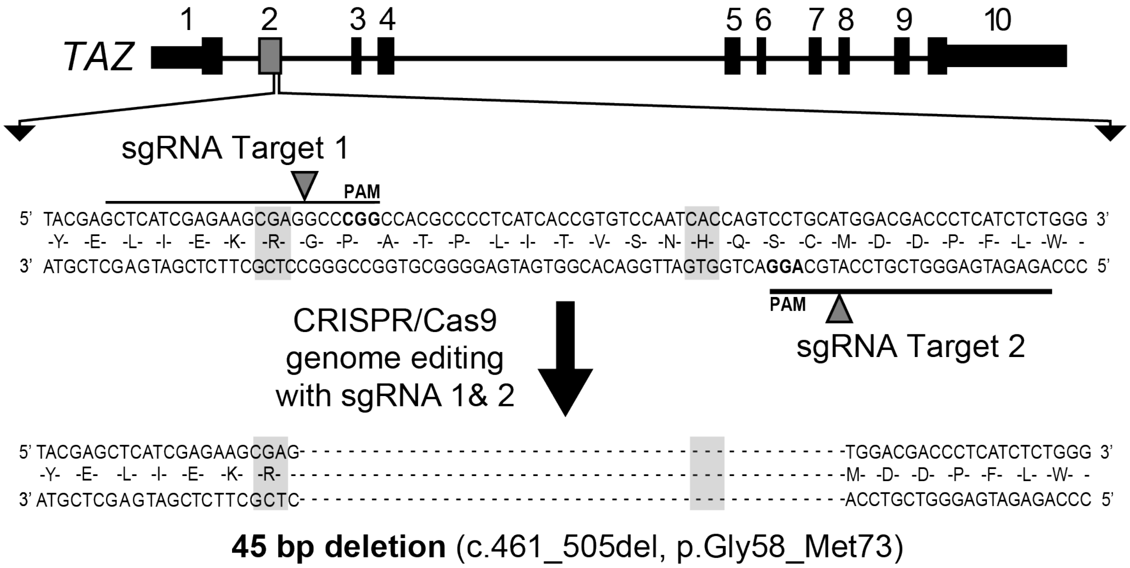


**Figure S1. CRISPR/Cas9 genome editing with two single guide RNAs (sgRNAs) targeting exon 2 of *TAZ* results in a 45 bp deletion.** The 45 bp deletion, noted by dashes, encompasses a predicted acyltransferase domain and covers an area of *TAZ* where multiple pathological variants have been described, such as p.R57L (c.170G>T) and p.H69Q (c.207C>G), highlighted in grey (3).

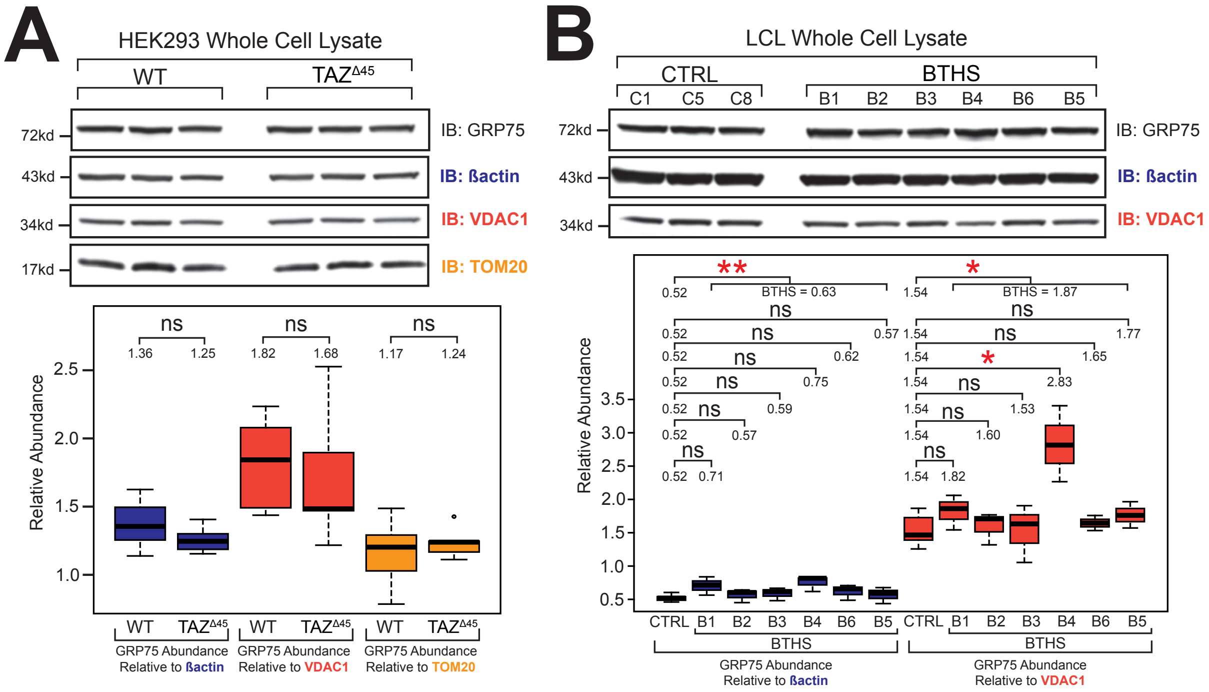


**Figure S2. Comparable abundance of cytosolic and mitochondrial proteins in the absence of TAZ.** Whole cell lysate (40 ug) of the indicated lines **(A)** HEK293 and **(B)** LCL lines were immunoblotted for the indicated proteins. Band intensities were quantified and plotted relative to ß-actin (blue), VDAC1 (red), and TOM20 (yellow); WT n=9, *TAZ*$^{\Delta45}$ n=9, CTRL n=9 (C1 n=3, C5 n=3, C8 n=3), BTHS n=18 (B1 n=3, B2 n=3, B3 n=3, B4 n=3, B6 n=3, B5 n=3). Significant differences are indicated; * ≤ 0.05, ** ≤ 0.005, *** ≤ 0.0005, **** ≤ 0.00005.

### Table S1. CRISPR/Cas9 sgRNA guide sequences and predicted off-target sites.

| sgRNA | Guide Sequence (5' to 3') | Off-Targets (5' to 3') | Score[1] | MMs[2] | Hg38 Location | Sequenced? |
|---|---|---|---|---|---|---|
| Target 1 | GCTCATCGAGAAGCGAGGCC**CGG**[3] | GCCCGTCAAGAAGCGAGGCCCAG | 2.4 | 3 | chr9:-136416984 | Y |
| | | GCTCATTGTTAAGCGAGGCCTAG | 0.9 | 3 | chr5:-164071618 | Y |
| | | CTTCAGCCAGAAGCGAGGCCAAG | 0.8 | 4 | chr13:+111688858 | Y |
| | | CCCCATCGAGAAGCGCGGCCAAG | 0.6 | 3 | chr4:-132956077 | N |
| | | GCCCATCGGGAAGCCAGGCCGAG | 0.5 | 3 | chr18:-8629183 | N |
| Target 2 | GAGATGAGGGTCGTCCATGC**AGG**[3] | GAGATGTAGCTCGTCCATGCTGG | 1.5 | 3 | chr3:+173436340 | Y |
| | | GTTCAGAGGGTCGTCCATGCAAG | 1.3 | 3 | chr7:-70923313 | Y |
| | | GATATGAGGGGAGTCCATGCAGG | 0.7 | 3 | chrX:-12667869 | N |
| | | GACAAGAGGTTGGTCCATGCCAG | 0.6 | 4 | chr7:+65692780 | N |
| | | CTGGTGAGGGTCTTCCATGCCAG | 0.6 | 4 | chr8:-125971306 | N |

[1] Off-target score calculated at cripsr.mit.edu based on scoring algorithm from Hsu et al. 2013 (2)
[2] The number of mismatches between the guide sequence and the "off-target" sequence
[3] PAM site is bolded

**Table S2A. Significant KEGG & GO terms determined by functional annotation analysis of the proteins with a FC ≤ 0.80 (n=215).**

| | # OF GENES[1] | %[2] | P-VALUE[3] | FOLD ENRICHMENT |
|---|---|---|---|---|
| **KEGG PATHWAYS** | | | | |
| Parkinson's disease | 13 | 6.1 | 6.80E-08 | 7.8 |
| Oxidative phosphorylation* | 11 | 5.1 | 2.70E-06 | 7 |
| Huntington's disease | 11 | 5.1 | 6.80E-05 | 4.9 |
| Alzheimer's disease | 10 | 4.7 | 1.30E-04 | 5.1 |
| Non-alcoholic fatty liver disease (NAFLD) | 9 | 4.2 | 3.40E-04 | 5.1 |
| Cardiac muscle contraction | 6 | 2.8 | 1.70E-03 | 6.8 |
| Bile secretion | 5 | 2.3 | 8.30E-03 | 6.2 |
| Dilated cardiomyopathy | 5 | 2.3 | 1.60E-02 | 5.1 |
| Metabolic pathways* | 23 | 10.7 | 1.90E-02 | 1.6 |
| Ribosome | 6 | 2.8 | 2.10E-02 | 3.7 |
| **GO TERMS: BIOLOGICAL PROCESSES** | | | | |
| platelet degranulation | 10 | 4.7 | 2.60E-06 | 8.5 |
| mitochondrial respiratory chain complex I assembly*# | 7 | 3.3 | 7.90E-05 | 9.7 |
| mitochondrial electron transport, NADH to ubiquinone*# | 6 | 2.8 | 2.30E-04 | 10.7 |
| actin filament organization | 6 | 2.8 | 1.40E-03 | 7.3 |
| response to oxidative stress | 7 | 3.3 | 1.60E-03 | 5.6 |
| protein lipoylation | 3 | 1.4 | 2.60E-03 | 37.5 |
| negative regulation of endothelial cell proliferation | 4 | 1.9 | 4.30E-03 | 12.1 |
| negative regulation of ATPase activity* | 3 | 1.4 | 5.50E-03 | 26.2 |
| aerobic respiration* | 4 | 1.9 | 6.10E-03 | 10.6 |
| protein targeting to mitochondrion* | 4 | 1.9 | 6.70E-03 | 10.3 |
| muscle contraction | 6 | 2.8 | 7.50E-03 | 4.9 |
| muscle filament sliding | 4 | 1.9 | 9.10E-03 | 9.2 |
| retina homeostasis | 4 | 1.9 | 1.10E-02 | 8.7 |
| rRNA processing | 8 | 3.7 | 1.10E-02 | 3.3 |
| wound healing | 5 | 2.3 | 1.30E-02 | 5.5 |
| Ossification | 5 | 2.3 | 1.30E-02 | 5.5 |
| cellular response to interferon-beta | 3 | 1.4 | 1.40E-02 | 16.4 |
| ribosomal small subunit biogenesis | 3 | 1.4 | 1.40E-02 | 16.4 |
| mitochondrial electron transport, cytochrome c to oxygen* | 3 | 1.4 | 2.10E-02 | 13.1 |
| SRP-dependent cotranslational protein targeting to membrane | 5 | 2.3 | 2.20E-02 | 4.7 |
| Translation | 8 | 3.7 | 2.60E-02 | 2.8 |
| cellular response to vascular endothelial growth factor stimulus | 3 | 1.4 | 2.80E-02 | 11.4 |
| response to calcium ion | 4 | 1.9 | 2.80E-02 | 6 |

| | | | | |
|---|---|---|---|---|
| response to electrical stimulus | 3 | 1.4 | 3.00E-02 | 10.9 |
| positive regulation of osteoblast differentiation | 4 | 1.9 | 3.10E-02 | 5.8 |
| cytoplasmic translation | 3 | 1.4 | 3.30E-02 | 10.5 |
| ribosomal large subunit biogenesis | 3 | 1.4 | 3.30E-02 | 10.5 |
| viral transcription | 5 | 2.3 | 3.90E-02 | 3.9 |
| positive regulation of phagocytosis | 3 | 1.4 | 4.30E-02 | 9 |
| sarcomere organization | 3 | 1.4 | 4.30E-02 | 9 |
| one-carbon metabolic process* | 3 | 1.4 | 4.50E-02 | 8.7 |
| nuclear-transcribed mRNA catabolic process, nonsense-mediated decay | 5 | 2.3 | 4.70E-02 | 3.7 |
| vascular endothelial growth factor receptor signaling pathway | 4 | 1.9 | 4.90E-02 | 4.9 |
| **GO TERMS: CELLULAR COMPARTMENTS** | | | | |
| mitochondrion* | 42 | 19.6 | 2.30E-09 | 2.8 |
| mitochondrial inner membrane* | 20 | 9.3 | 5.80E-07 | 4 |
| extracellular exosome | 55 | 25.7 | 3.10E-05 | 1.7 |
| prefoldin complex | 4 | 1.9 | 4.70E-05 | 50.8 |
| mitochondrial respiratory chain complex I*# | 6 | 2.8 | 2.10E-04 | 10.9 |
| platelet alpha granule lumen | 6 | 2.8 | 3.70E-04 | 9.7 |
| Cytosol | 57 | 26.6 | 6.90E-04 | 1.5 |
| Cytoskeleton | 13 | 6.1 | 1.00E-03 | 3.1 |
| cell surface | 16 | 7.5 | 1.20E-03 | 2.6 |
| focal adhesion | 13 | 6.1 | 1.60E-03 | 3 |
| mitochondrial proton-transporting ATP synthase complex* | 4 | 1.9 | 1.60E-03 | 16.9 |
| mitochondrial membrane* | 6 | 2.8 | 4.10E-03 | 5.7 |
| actin filament | 5 | 2.3 | 6.10E-03 | 6.8 |
| cytosolic large ribosomal subunit | 5 | 2.3 | 7.10E-03 | 6.5 |
| basement membrane | 5 | 2.3 | 1.20E-02 | 5.6 |
| Costamere | 3 | 1.4 | 1.90E-02 | 14 |
| integral component of mitochondrial inner membrane* | 3 | 1.4 | 2.10E-02 | 13.3 |
| respiratory chain* | 3 | 1.4 | 2.10E-02 | 13.3 |
| stress fiber | 4 | 1.9 | 2.30E-02 | 6.6 |
| Microvillus | 4 | 1.9 | 2.60E-02 | 6.2 |
| M band | 3 | 1.4 | 2.70E-02 | 11.6 |
| blood microparticle | 6 | 2.8 | 2.80E-02 | 3.5 |
| myelin sheath | 6 | 2.8 | 2.80E-02 | 3.5 |
| Cytoplasm | 72 | 33.6 | 3.20E-02 | 1.2 |
| Microspike | 2 | 0.9 | 3.30E-02 | 59.3 |
| granular component | 2 | 0.9 | 4.40E-02 | 44.4 |
| muscle thin filament tropomyosin | 2 | 0.9 | 4.40E-02 | 44.4 |
| apical plasma membrane | 8 | 3.7 | 4.60E-02 | 2.4 |

| | | | | |
|---|---|---|---|---|
| filamentous actin | 3 | 1.4 | 4.70E-02 | 8.6 |
| extracellular matrix | 8 | 3.7 | 4.90E-02 | 2.4 |
| **GO TERMS: MOLECULAR FUNCTIONS** | | | | |
| structural constituent of muscle | 6 | 2.8 | 1.10E-04 | 12.6 |
| protein binding | 125 | 58.4 | 2.00E-04 | 1.3 |
| actin filament binding | 8 | 3.7 | 7.60E-04 | 5.3 |
| unfolded protein binding | 7 | 3.3 | 1.60E-03 | 5.6 |
| NADH dehydrogenase (ubiquinone) activity *# | 5 | 2.3 | 2.10E-03 | 9.2 |
| poly(A) RNA binding | 25 | 11.7 | 2.10E-03 | 1.9 |
| structural constituent of ribosome | 9 | 4.2 | 3.80E-03 | 3.6 |
| endopeptidase inhibitor activity | 4 | 1.9 | 1.00E-02 | 8.8 |
| identical protein binding | 17 | 7.9 | 1.10E-02 | 2 |
| structural molecule activity conferring elasticity | 2 | 0.9 | 2.30E-02 | 87.9 |
| structural constituent of cytoskeleton | 5 | 2.3 | 3.60E-02 | 4 |
| ubiquitin conjugating enzyme activity | 3 | 1.4 | 4.20E-02 | 9.1 |
| cytochrome-c oxidase activity * | 3 | 1.4 | 4.50E-02 | 8.8 |
| * References mitochondrion and/or metabolic pathways (n=18)<br># References complex I of the oxidative phosphorylation pathways (n=4) | | | | |
| [1] The number of input proteins involved in the term<br>[2] The number of input proteins involved in the term divided by the total proteins/genes represented by the term<br>[3] Modified Fisher Exact p-value, EASE Score | | | | |

**Table S2B. Significant KEGG & GO terms determined by functional annotation analysis of the proteins with a FC ≥ 1.20 (n=621).**

| | # OF GENES[1] | %[2] | P-VALUE[3] | FOLD ENRICHMENT |
|---|---|---|---|---|
| **KEGG PATHWAYS** | | | | |
| Chronic myeloid leukemia | 9 | 1.4 | 2.50E-03 | 3.8 |
| Metabolic pathways* | 58 | 9.3 | 3.10E-03 | 1.4 |
| Pancreatic cancer | 8 | 1.3 | 5.40E-03 | 3.7 |
| AMPK signaling pathway | 11 | 1.8 | 7.50E-03 | 2.7 |
| Alzheimer’s disease | 13 | 2.1 | 9.70E-03 | 2.3 |
| Butanoate metabolism* | 5 | 0.8 | 1.10E-02 | 5.6 |
| Carbohydrate digestion and absorption | 6 | 1 | 1.20E-02 | 4.3 |
| Endocytosis | 16 | 2.6 | 1.40E-02 | 2 |
| Insulin signaling pathway | 11 | 1.8 | 1.60E-02 | 2.4 |
| Colorectal cancer | 7 | 1.1 | 1.60E-02 | 3.4 |
| Fatty acid metabolism* | 6 | 1 | 2.10E-02 | 3.8 |
| Notch signaling pathway | 6 | 1 | 2.10E-02 | 3.8 |

| | | | | |
|---|---|---|---|---|
| Amino sugar and nucleotide sugar metabolism* | 6 | 1 | 2.10E-02 | 3.8 |
| p53 signaling pathway | 7 | 1.1 | 2.30E-02 | 3.1 |
| Biosynthesis of antibiotics | 14 | 2.3 | 2.30E-02 | 2 |
| Phagosome | 11 | 1.8 | 2.70E-02 | 2.2 |
| Adipocytokine signaling pathway | 7 | 1.1 | 2.80E-02 | 3 |
| Inositol phosphate metabolism* | 7 | 1.1 | 3.00E-02 | 3 |
| Proteoglycans in cancer | 13 | 2.1 | 3.30E-02 | 2 |
| Biosynthesis of unsaturated fatty acids | 4 | 0.6 | 3.90E-02 | 5.2 |
| Viral myocarditis | 6 | 1 | 4.00E-02 | 3.2 |
| Synthesis and degradation of ketone bodies | 3 | 0.5 | 4.10E-02 | 9 |
| Phosphatidylinositol signaling system | 8 | 1.3 | 4.30E-02 | 2.5 |
| Glucagon signaling pathway | 8 | 1.3 | 4.60E-02 | 2.4 |
| Cell adhesion molecules (CAMs) | 10 | 1.6 | 4.60E-02 | 2.1 |
| Fatty acid elongation | 4 | 0.6 | 4.80E-02 | 4.8 |
| Epstein-Barr virus infection | 9 | 1.4 | 4.90E-02 | 2.2 |
| **GO TERMS: BIOLOGICAL PROCESSES** | | | | |
| covalent chromatin modification | 14 | 2.3 | 7.30E-05 | 3.8 |
| viral genome replication | 5 | 0.8 | 8.40E-04 | 11 |
| protein targeting to plasma membrane | 6 | 1 | 1.30E-03 | 7.1 |
| antigen processing and presentation of endogenous peptide antigen via MHC class I via ER pathway, TAP-independent | 3 | 0.5 | 3.10E-03 | 30.9 |
| positive regulation of apoptotic process* | 20 | 3.2 | 4.20E-03 | 2.1 |
| carbohydrate phosphorylation | 5 | 0.8 | 5.90E-03 | 6.7 |
| cell cycle arrest | 12 | 1.9 | 6.10E-03 | 2.6 |
| response to oxidative stress* | 10 | 1.6 | 9.30E-03 | 2.8 |
| cell migration | 13 | 2.1 | 1.00E-02 | 2.3 |
| phosphatidylinositol biosynthetic process | 7 | 1.1 | 1.10E-02 | 3.7 |
| long-chain fatty-acyl-CoA biosynthetic process | 6 | 1 | 1.10E-02 | 4.4 |
| macroautophagy* | 8 | 1.3 | 1.10E-02 | 3.2 |
| unsaturated fatty acid biosynthetic process | 4 | 0.6 | 1.10E-02 | 8.2 |
| muscle cell differentiation | 4 | 0.6 | 1.10E-02 | 8.2 |
| IRE1-mediated unfolded protein response | 7 | 1.1 | 1.20E-02 | 3.7 |
| positive regulation of DNA binding | 5 | 0.8 | 1.20E-02 | 5.5 |
| tRNA pseudouridine synthesis | 3 | 0.5 | 1.40E-02 | 15.4 |
| magnesium ion homeostasis | 3 | 0.5 | 1.40E-02 | 15.4 |
| antigen processing and presentation of peptide antigen via MHC class I | 5 | 0.8 | 1.50E-02 | 5.1 |
| neuron projection development | 9 | 1.4 | 1.60E-02 | 2.8 |
| positive regulation of substrate adhesion-dependent cell spreading | 5 | 0.8 | 1.90E-02 | 4.8 |
| membrane protein intracellular domain proteolysis | 4 | 0.6 | 1.90E-02 | 6.9 |
| cilium assembly | 10 | 1.6 | 1.90E-02 | 2.5 |

| | | | | |
|---|---|---|---|---|
| regulation of apoptotic process* | 14 | 2.3 | 2.10E-02 | 2 |
| inner ear receptor stereocilium organization | 4 | 0.6 | 2.20E-02 | 6.5 |
| regulation of autophagy* | 6 | 1 | 2.20E-02 | 3.7 |
| amyloid precursor protein catabolic process | 3 | 0.5 | 2.60E-02 | 11.6 |
| response to cholesterol | 3 | 0.5 | 2.60E-02 | 11.6 |
| interferon-gamma-mediated signaling pathway | 7 | 1.1 | 2.70E-02 | 3 |
| protein maturation by protein folding | 3 | 0.5 | 3.20E-02 | 10.3 |
| pyrimidine nucleotide metabolic process* | 3 | 0.5 | 3.20E-02 | 10.3 |
| DNA topological change | 3 | 0.5 | 3.20E-02 | 10.3 |
| antigen processing and presentation of exogenous peptide antigen via MHC class I, TAP-independent | 3 | 0.5 | 3.20E-02 | 10.3 |
| cilium morphogenesis | 10 | 1.6 | 3.30E-02 | 2.3 |
| cell growth | 6 | 1 | 3.40E-02 | 3.3 |
| viral process | 17 | 2.7 | 3.40E-02 | 1.8 |
| purine nucleotide metabolic process* | 3 | 0.5 | 4.00E-02 | 9.3 |
| regulation of synaptic vesicle exocytosis | 3 | 0.5 | 4.00E-02 | 9.3 |
| cell volume homeostasis | 3 | 0.5 | 4.00E-02 | 9.3 |
| nucleocytoplasmic transport | 4 | 0.6 | 4.60E-02 | 4.9 |
| positive regulation of stress fiber assembly* | 5 | 0.8 | 4.60E-02 | 3.7 |
| UDP-N-acetylglucosamine biosynthetic process | 3 | 0.5 | 4.70E-02 | 8.4 |
| positive regulation of histone deacetylation | 3 | 0.5 | 4.70E-02 | 8.4 |
| chromatin organization | 5 | 0.8 | 4.90E-02 | 3.6 |
| histone H3 acetylation | 5 | 0.8 | 4.90E-02 | 3.6 |
| glycosaminoglycan catabolic process | 4 | 0.6 | 5.00E-02 | 4.7 |
| **GO TERMS: CELLULAR COMPARTMENTS** | | | | |
| extracellular exosome | 135 | 21.7 | 1.70E-06 | 1.5 |
| membrane | 110 | 17.7 | 3.70E-06 | 1.5 |
| endoplasmic reticulum | 51 | 8.2 | 1.90E-05 | 1.9 |
| mitochondrion* | 68 | 11 | 2.20E-04 | 1.6 |
| cytosol | 142 | 22.9 | 2.80E-04 | 1.3 |
| cytoplasm | 206 | 33.2 | 7.60E-04 | 1.2 |
| nucleoplasm | 120 | 19.3 | 8.10E-04 | 1.3 |
| integral component of endoplasmic reticulum membrane | 11 | 1.8 | 2.00E-03 | 3.3 |
| integral component of luminal side of endoplasmic reticulum membrane | 6 | 1 | 2.20E-03 | 6.4 |
| phagocytic vesicle membrane | 8 | 1.3 | 2.90E-03 | 4.2 |
| endoplasmic reticulum membrane | 44 | 7.1 | 3.60E-03 | 1.6 |
| intermediate filament | 11 | 1.8 | 3.80E-03 | 3 |
| ruffle membrane | 9 | 1.4 | 5.10E-03 | 3.4 |
| mitochondrial inner membrane* | 26 | 4.2 | 5.20E-03 | 1.8 |
| ER to Golgi transport vesicle membrane | 7 | 1.1 | 6.60E-03 | 4.1 |

| | | | | |
|---|---|---|---|---|
| mitochondrial intermembrane space* | 8 | 1.3 | 1.00E-02 | 3.3 |
| myelin sheath | 12 | 1.9 | 1.10E-02 | 2.4 |
| nucleus | 202 | 32.5 | 1.30E-02 | 1.1 |
| melanosome | 9 | 1.4 | 1.70E-02 | 2.7 |
| early endosome | 15 | 2.4 | 1.80E-02 | 2 |
| Golgi membrane | 30 | 4.8 | 1.80E-02 | 1.6 |
| centrosome | 23 | 3.7 | 2.20E-02 | 1.7 |
| costamere | 4 | 0.6 | 2.20E-02 | 6.5 |
| endoplasmic reticulum-Golgi intermediate compartment | 7 | 1.1 | 2.30E-02 | 3.2 |
| endoplasmic reticulum lumen | 13 | 2.1 | 2.30E-02 | 2.1 |
| cilium | 11 | 1.8 | 2.60E-02 | 2.2 |
| autolysosome | 3 | 0.5 | 2.60E-02 | 11.5 |
| ER-mitochondrion membrane contact site* | 3 | 0.5 | 2.60E-02 | 11.5 |
| integral component of mitochondrial outer membrane* | 4 | 0.6 | 2.90E-02 | 5.9 |
| stress fiber* | 6 | 1 | 3.00E-02 | 3.4 |
| F-actin capping protein complex | 3 | 0.5 | 3.30E-02 | 10.2 |
| nuclear heterochromatin | 4 | 0.6 | 3.30E-02 | 5.6 |
| lysosomal membrane | 16 | 2.6 | 3.50E-02 | 1.8 |
| early endosome membrane | 9 | 1.4 | 3.50E-02 | 2.4 |
| Golgi apparatus | 39 | 6.3 | 3.60E-02 | 1.4 |
| oligosaccharyltransferase complex | 3 | 0.5 | 4.00E-02 | 9.2 |
| dendrite | 18 | 2.9 | 4.70E-02 | 1.7 |
| MHC class I protein complex | 3 | 0.5 | 4.80E-02 | 8.4 |
| **GO TERMS: MOLECULAR FUNCTIONS** | | | | |
| protein binding | 335 | 53.9 | 2.90E-05 | 1.2 |
| poly(A) RNA binding | 55 | 8.9 | 3.10E-03 | 1.5 |
| AP-3 adaptor complex binding | 3 | 0.5 | 3.10E-03 | 30.5 |
| TAP binding | 3 | 0.5 | 3.10E-03 | 30.5 |
| nucleosomal DNA binding | 7 | 1.1 | 3.70E-03 | 4.6 |
| cysteine-type endopeptidase activity involved in apoptotic process* | 4 | 0.6 | 7.80E-03 | 9.4 |
| chromatin binding | 23 | 3.7 | 9.80E-03 | 1.8 |
| 1-phosphatidylinositol-3-phosphate 4-kinase activity | 3 | 0.5 | 1.00E-02 | 18.3 |
| GTPase activity | 16 | 2.6 | 1.00E-02 | 2.1 |
| peptide antigen binding | 5 | 0.8 | 1.20E-02 | 5.5 |
| GTP binding | 22 | 3.5 | 1.50E-02 | 1.7 |
| histone binding | 10 | 1.6 | 1.90E-02 | 2.5 |
| scaffold protein binding | 6 | 1 | 2.00E-02 | 3.8 |
| 1-phosphatidylinositol-4-phosphate 5-kinase activity | 3 | 0.5 | 2.00E-02 | 13.1 |
| chromatin DNA binding | 6 | 1 | 4.10E-02 | 3.2 |

| | | | | |
|---|---|---|---|---|
| cysteine-type endopeptidase activity | 6 | 1 | 4.90E-02 | 3 |
| * References mitochondrion and/or mitochondrial dynamics (n=20) | | | | |
| [1] The number of input proteins involved in the term<br>[2] The number of input proteins involved in the term divided by the total proteins/genes represented by the term<br>[3] Modified Fisher Exact p-value, EASE Score | | | | |

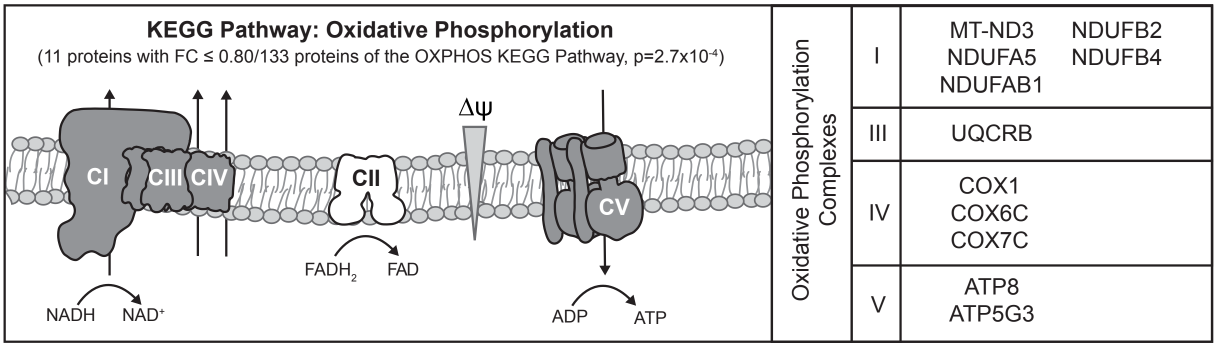


**Figure S3. The oxidative phosphorylation (OXPHOS) KEGG pathway is the most significant KEGG pathway enriched for proteins with a FC ≤ 0.80 that references mitochondria and or OXPHOS.** Of the 133 genes in the OXPHOS KEGG pathway 11 encode proteins with a FC ≤ 0.80 in *TAZ*$^{\Delta 45}$ cells; half (n=5) are subunits of complex I (CI) and the remaining (n=6) are subunits of complex III, IV, and V.

### Table S3. Proteomics quantification of all complex I (CI) associated proteins.

| UniProt Entry | Gene Name | Module | Protein | #PSMs* | Unique Peptides | FC | P-Value |
|---|---|---|---|---|---|---|---|
| **Complex I Subunits** | | | | | | | |
| Q9UI09 | *NDUFA12* | N | NADH dehydrogenase [ubiquinone] 1 alpha subcomplex subunit 12 | 46 | 8 | 0.973 | 0.93 |
| P28331 | *NDUFS1* | N | NADH-ubiquinone oxidoreductase 75 kDa subunit, mitochondrial | 173 | 39 | 0.864 | 0.28 |
| O75380 | *NDUFS6* | N | NADH dehydrogenase [ubiquinone] iron-sulfur protein 6, mitochondrial | 23 | 9 | 0.955 | 0.49 |
| P49821 | *NDUFV1* | N | NADH dehydrogenase [ubiquinone] flavoprotein 1, mitochondrial | 105 | 21 | 0.872 | 0.21 |
| P19404 | *NDUFV2* | N | NADH dehydrogenase [ubiquinone] flavoprotein 2, mitochondrial | 73 | 15 | 0.872 | 0.18 |
| P56181 | *NDUFV3* | N | NADH dehydrogenase [ubiquinone] flavoprotein 3, mitochondrial | 5 | 3 | 1.037 | 0.69 |
| O43678 | *NDUFA2* | N/Q | NADH dehydrogenase [ubiquinone] 1 alpha subcomplex subunit 2 | 19 | 4 | 0.862 | 0.15 |
| O43181 | *NDUFS4* | N/Q | NADH dehydrogenase [ubiquinone] iron-sulfur protein 4, mitochondrial | 21 | 7 | 0.890 | 0.22 |
| **Q16718** | ***NDUFA5*** | **Q** | **NADH dehydrogenase [ubiquinone] 1 alpha subcomplex subunit 5** | **33** | **7** | **0.722** | **0.24** |
| P56556 | *NDUFA6* | Q | NADH dehydrogenase [ubiquinone] 1 alpha subcomplex subunit 6 | 27 | 7 | 0.902 | 0.89 |
| O95182 | *NDUFA7* | Q | NADH dehydrogenase [ubiquinone] 1 alpha subcomplex subunit 7 | 33 | 9 | 0.834 | 0.21 |
| Q16795 | *NDUFA9* | Q | NADH dehydrogenase [ubiquinone] 1 alpha subcomplex subunit 9, mitochondrial | 54 | 18 | 0.970 | 0.99 |
| O75306 | *NDUFS2* | Q | NADH dehydrogenase [ubiquinone] iron-sulfur protein 2, mitochondrial | 130 | 22 | 1.018 | 0.95 |
| O75489 | *NDUFS3* | Q | NADH dehydrogenase [ubiquinone] iron-sulfur protein 3, mitochondrial | 142 | 18 | 0.953 | 0.53 |
| O75251 | *NDUFS7* | Q | NADH dehydrogenase [ubiquinone] iron-sulfur protein 7, mitochondrial | 27 | 6 | 0.873 | 0.94 |
| O00217 | *NDUFS8* | Q | NADH dehydrogenase [ubiquinone] iron-sulfur protein 8, mitochondrial | 66 | 9 | 0.959 | 0.46 |
| P03886 | *MTND1* | $P_P$ | NADH-ubiquinone oxidoreductase chain 1 | 4 | 2 | 1.236 | 0.43 |
| P03891 | *MTND2* | $P_P$ | NADH-ubiquinone oxidoreductase chain 2 | 3 | 2 | 1.358 | 0.24 |
| **P03897** | ***MTND3**** | $\mathbf{P_P}$ | **NADH-ubiquinone oxidoreductase chain 3** | **2** | **1** | **0.608** | **0.01** |
| P03923 | *MTND6* | $P_P$ | NADH-ubiquinone oxidoreductase chain 6 | 2 | 1 | 1.391 | 0.45 |
| O95299 | *NDUFA10* | $P_P$ | NADH dehydrogenase [ubiquinone] 1 alpha subcomplex subunit 10, mitochondrial | 56 | 17 | 0.957 | 0.82 |
| Q86Y39 | *NDUFA11* | $P_P$ | NADH dehydrogenase [ubiquinone] 1 alpha subcomplex subunit 11 | 14 | 3 | 0.84 | 0.20 |
| Q9P0J0 | *NDUFA13* | $P_P$ | NADH dehydrogenase [ubiquinone] 1 alpha subcomplex subunit 13 | 20 | 8 | 0.986 | 0.90 |
| O95167 | *NDUFA3* | $P_P$ | NADH dehydrogenase [ubiquinone] 1 alpha subcomplex subunit 3 | 17 | 4 | 0.96 | 0.97 |
| P51970 | *NDUFA8* | $P_P$ | NADH dehydrogenase [ubiquinone] 1 alpha subcomplex subunit 8 | 31 | 7 | 0.937 | 0.43 |
| O95298 | *NDUFC2* | $P_P$ | NADH dehydrogenase [ubiquinone] 1 subunit C2 | 19 | 6 | 0.808 | 0.10 |
| O43920 | *NDUFS5* | $P_P$ | NADH dehydrogenase [ubiquinone] iron-sulfur protein 5 | 29 | 7 | 0.935 | 0.22 |
| P03905 | *MTND4* | $P_D$ | NADH-ubiquinone oxidoreductase chain 4 | 3 | 2 | 1.036 | 0.57 |
| P03915 | *MTND5* | $P_D$ | NADH-ubiquinone oxidoreductase chain 5 | 11 | 3 | 0.977 | 1.00 |
| **O14561** | ***NDUFAB1*** | $\mathbf{P_D}$ | **Acyl carrier protein, mitochondrial** | **6** | **2** | **0.727** | **0.46** |
| O75438 | *NDUFB1* | $P_D$ | NADH dehydrogenase [ubiquinone] 1 beta subcomplex subunit 1 | 3 | 2 | 0.995 | 0.98 |
| O96000 | *NDUFB10* | $P_D$ | NADH dehydrogenase [ubiquinone] 1 beta subcomplex subunit 10 | 40 | 8 | 0.904 | 0.47 |
| Q9NX14 | *NDUFB11* | $P_D$ | NADH dehydrogenase [ubiquinone] 1 beta subcomplex subunit 11, mitochondrial | 20 | 3 | 0.863 | 0.10 |
| **O95178** | ***NDUFB2*** | $\mathbf{P_D}$ | **NADH dehydrogenase [ubiquinone] 1 beta subcomplex subunit 2** | **6** | **1** | **0.62** | **0.12** |
| O43676 | *NDUFB3*** | $P_D$ | NADH dehydrogenase [ubiquinone] 1 beta subcomplex subunit 3 | 11 | 3 | 0.864 | 0.05 |
| **O95168** | ***NDUFB4*** | $\mathbf{P_D}$ | **NADH dehydrogenase [ubiquinone] 1 beta subcomplex subunit 4** | **30** | **4** | **0.762** | **0.22** |
| O43674 | *NDUFB5* | $P_D$ | NADH dehydrogenase [ubiquinone] 1 beta subcomplex subunit 5, mitochondrial | 28 | 6 | 0.918 | 1.00 |
| O95139 | *NDUFB6* | $P_D$ | NADH dehydrogenase [ubiquinone] 1 beta subcomplex subunit 6 | 18 | 7 | 1.081 | 0.99 |
| P17568 | *NDUFB7* | $P_D$ | NADH dehydrogenase [ubiquinone] 1 beta subcomplex subunit 7 | 11 | 4 | 0.814 | 0.30 |
| O95169 | *NDUFB8* | $P_D$ | NADH dehydrogenase [ubiquinone] 1 beta subcomplex subunit 8, mitochondrial | 13 | 4 | 0.911 | 0.25 |
| Q9Y6M9 | *NDUFB9* | $P_D$ | NADH dehydrogenase [ubiquinone] 1 beta subcomplex subunit 9 | 38 | 8 | 0.92 | 0.17 |
| **Complex I Assembly Factors** | | | | | | | |
| Q9H845 | *ACAD9* | | Acyl-CoA dehydrogenase family member 9, mitochondrial | 80 | 30 | 1.107 | 0.15 |
| Q9GZY4 | *COA1* | | Cytochrome c oxidase assembly factor 1 homolog | 6 | 4 | 1.119 | 0.78 |
| Q9BQ95 | *ECSIT* | | Evolutionarily conserved signaling intermediate in Toll pathway, mitochondrial | 12 | 9 | 0.918 | 0.47 |
| Q96CU9 | *FOXRED1* | | FAD-dependent oxidoreductase domain-containing protein 1 | 13 | 6 | 0.96 | 0.97 |
| **Q9Y375** | ***NDUFAF1**** | | **Complex I intermediate-associated protein 30, mitochondrial** | **11** | **8** | **0.797** | **0.004** |
| Q8N183 | *NDUFAF2* | | NADH dehydrogenase [ubiquinone] 1 alpha subcomplex assembly factor 2 | 49 | 13 | 0.855 | 0.35 |
| Q9BU61 | *NDUFAF3* | | NADH dehydrogenase [ubiquinone] 1 alpha subcomplex assembly factor 3 | 32 | 7 | 0.942 | 0.88 |
| Q9P032 | *NDUFAF4* | | NADH dehydrogenase [ubiquinone] 1 alpha subcomplex assembly factor 4 | 30 | 10 | 0.952 | 0.30 |
| Q5TEU4 | *NDUFAF5* | | Arginine-hydroxylase NDUFAF5, mitochondrial | 1 | 1 | 0.98 | 0.73 |
| Q330K2 | *NDUFAF6* | | NADH dehydrogenase (ubiquinone) complex I, assembly factor 6 | 1 | 1 | 0.998 | 0.86 |

| | | | | | | |
|---|---|---|---|---|---|---|
| Q7L592 | *NDUFAF7* | Protein arginine methyltransferase NDUFAF7, mitochondrial | 16 | 10 | 0.95 | 0.99 |
| Q8TB37 | *NUBPL* | Iron-sulfur protein NUBPL | 7 | 5 | 0.952 | 1.00 |
| Q9NPL8 | *TIMMDC1* | Complex I assembly factor TIMMDC1, mitochondrial | 23 | 8 | 1.118 | 0.02 |
| Q8IUX1 | *TMEM126B* | Complex I assembly factor TMEM126B, mitochondrial | 1 | 1 | 0.984 | 0.96 |
| Q9BUB7 | *TMEM70* | Transmembrane protein 70, mitochondrial | 5 | 3 | 1.063 | 0.84 |

[*] Peptide spectral matches
[Bold Font] Subunits or assembly factors with FC ≤ 0.80 (n=6)
[**] Significantly reduced subunits or assembly factors (n=2)

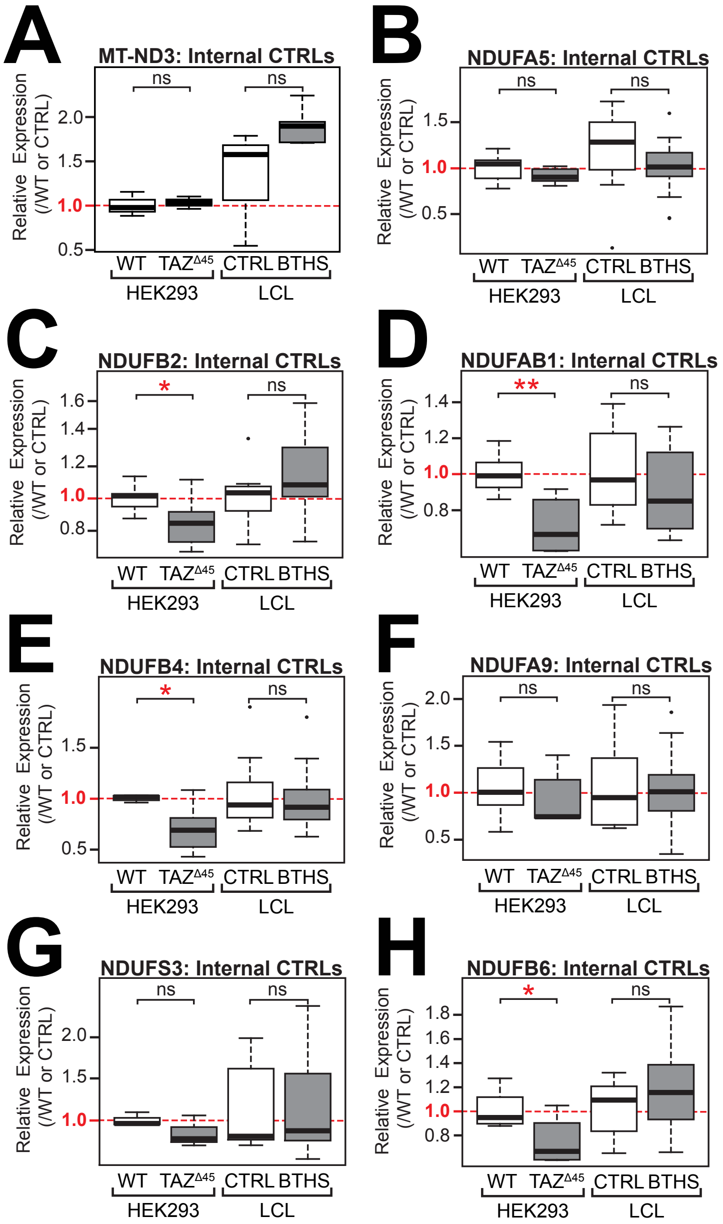


**Figure S4. Relative mRNA expression of complex I (CI) subunits; (A)** *MT-ND3* **(B)** *NDUFA5* **(C)** *NDUFB2* **(D)** *NDUFAB1* **(E)** *NDUFB4* **(F)** *NDUFA9* **(G)** *NDUFS3* **(H)** *NDUFB6* determined by qRT-PCR and $\Delta\Delta C_T$ quantification; WT n=6 (except *MT-ND3* and *NDUFS3*, n=3), $TAZ^{\Delta 45}$ n=6 (except *MT-ND3* and *NDUFS3*, n=3), CTRL n=10 (except *MT-ND3* n=5), BTHS n=15 (except *MT-ND3* n=3). Significant differences are indicated; * ≤ 0.05, ** ≤ 0.005, *** ≤ 0.0005, **** ≤ 0.00005.

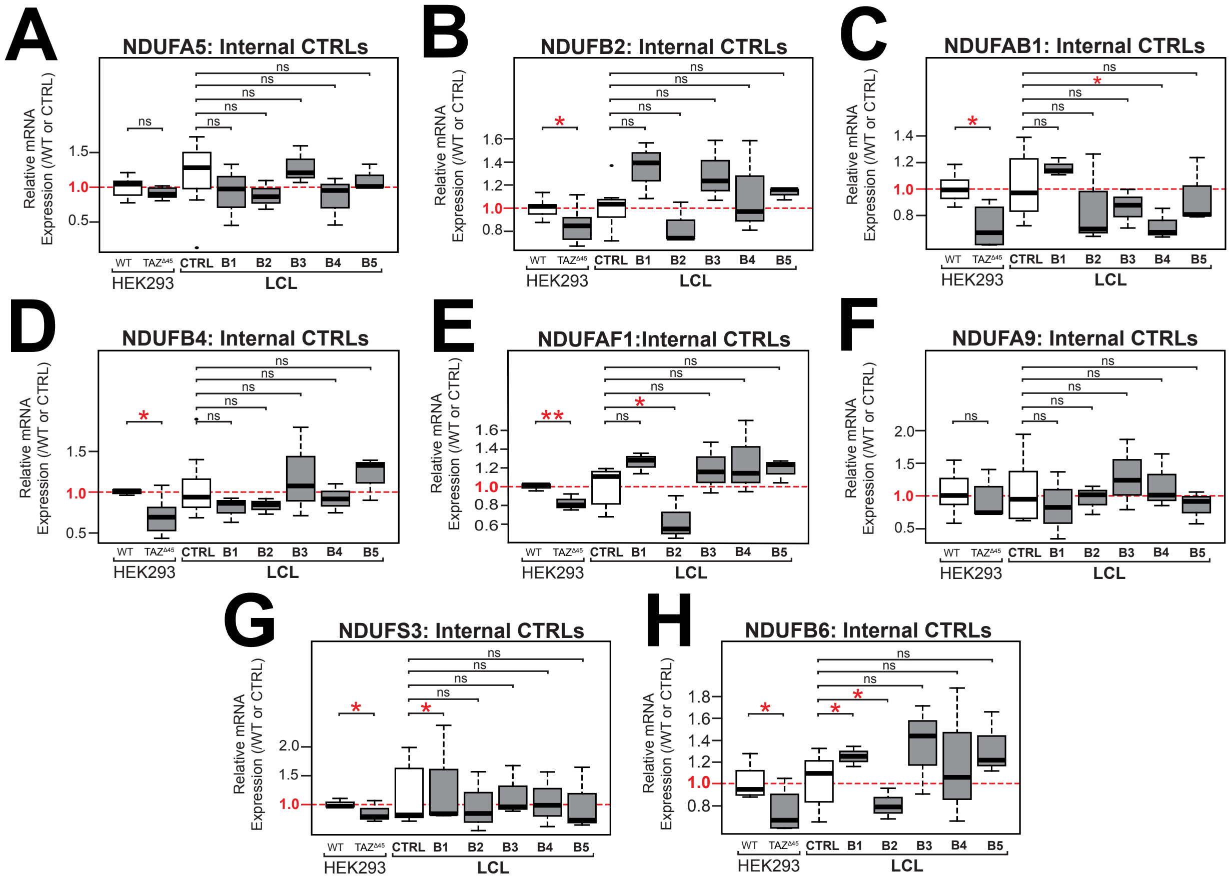


**Figure S5. Relative mRNA expression of complex I (CI) subunits varies across different LCL lines.** Relative mRNA expression of **(A)** *NDUFA5* **(B)** *NDUFB2* **(C)** *NDUFAB1* **(D)** *NDUFB4* **(E)** *NDUFAF1* **(F)** *NDUFA9* **(G)** *NDUFS3* **(H)** *NDUFB6* determined by qRT-PCR and $\Delta\Delta C_T$ quantification; WT n=6 (except *MT-ND3* and *NDUFS3*, n=3), *TAZ*$^{\Delta 45}$ n=6 (except *MT-ND3* and *NDUFS3*, n=3), CTRL n=10, B1 n=3, B2 n=3, B3 n=3, B4 n=3, B5 n=3. Significant differences are indicated; * ≤ 0.05, ** ≤ 0.005, *** ≤ 0.0005, **** ≤ 0.00005.

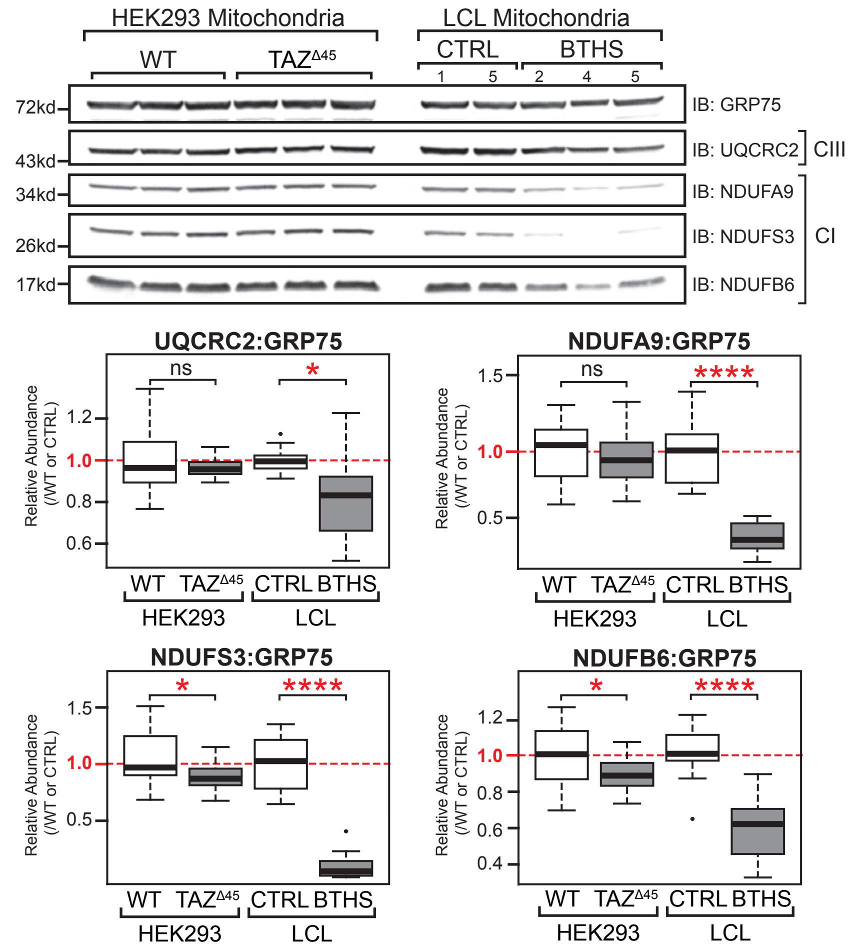


**Figure S6. Immunoblotting of isolated mitochondria for CI and CIII subunits.** Mitochondria (40 ug) isolated from the indicated lines were immunoblotted for the indicated proteins. Band intensities, relative to loading control GRP75, were quantified and plotted relative to WT/CTRL abundance; WT n=15, $TAZ^{\Delta 45}$ n=15, CTRL n=10, BTHS n=15. Significant differences are indicated; * ≤ 0.05, ** ≤ 0.005, *** ≤ 0.0005, **** ≤ 0.00005.

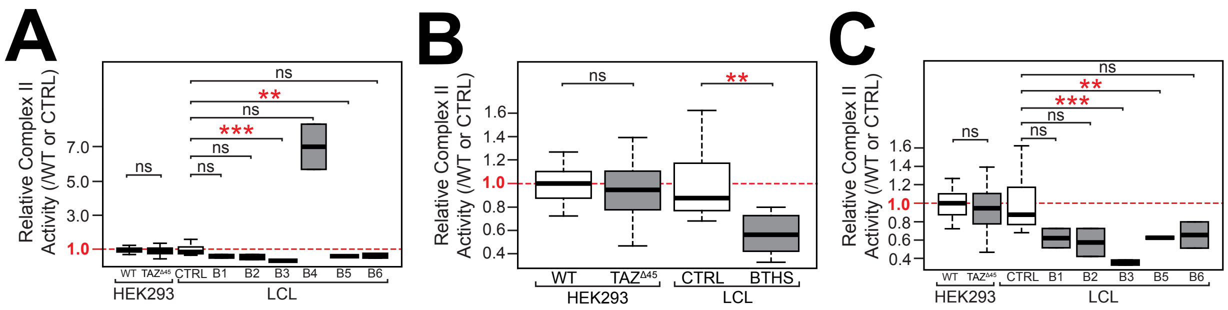


**Figure S7. Difference in complex II (CII) activity between and within cell types.** CII activity measured in HEK293 WT and *TAZ*$^{\Delta45}$ or CTRL and BTHS LCLs mitochondria (200 ug total protein) on a microplate reader by following the production of ubiquinol by CII coupled to the reduction of the dye diclorophenolindophenol (DCPIP, 600nm). **(A)** Analysis of CII activity in LCLs showed that a single BTHS line, BTHS LCL #4, had significantly increased CII activity compared to the other BTHS lines; WT n=12, *TAZ*$^{\Delta45}$ n=12, CTRL n=10, B1 n=2, B2 n=2, B3 n=2, B4 n=2, B5 n=2, B6 n=2. **(B)** Activity plotted without outlier BTHS LCL #4; WT n=12, *TAZ*$^{\Delta45}$ n=12, CTRL n=10, BTHS n=10. **(C)** WT n=12, *TAZ*$^{\Delta45}$ n=12, CTRL n=10, B1 n=2, B2 n=2, B3 n=2, B5 n=2, B6 n=2. Significant differences are indicated; * ≤ 0.05, ** ≤ 0.005, *** ≤ 0.0005, **** ≤ 0.00005.

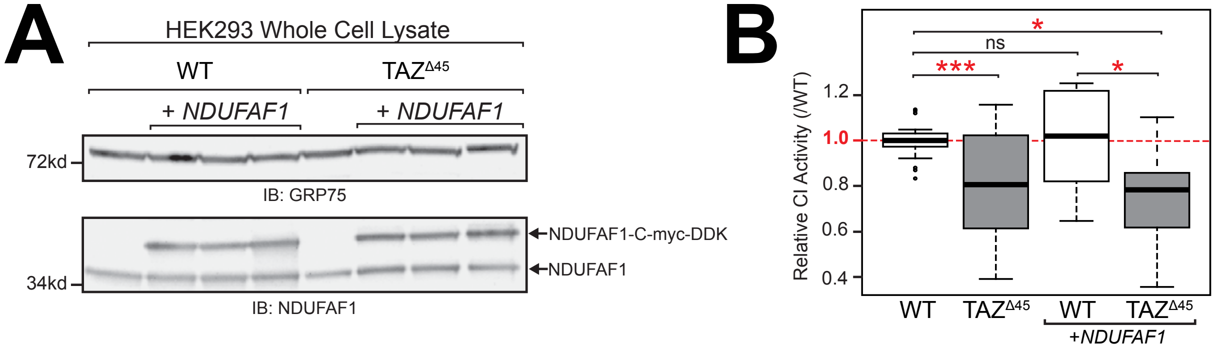


**Figure S8. Overexpression of CI assembly factor NDUFAF1 does not normalize CI activity.** **(A)** HEK293 WT and *TAZ*$^{\Delta 45}$ cells were transiently transfected with tagged *NDUFAF1* with Lipofectamine 3000 according to manufacturer's instructions. Whole cell extracts (45 ug) of the indicated lines and treatment concentrations were immunoblotted for the indicated proteins. **(B)** CI activity measured in mitochondria (200 ug total protein). Activity was measured on a microplate reader (450nm) by following the oxidation of NADH to oxidized nicotinamide adenine dinucleotide (NAD+). Activity plotted relative to WT abundance; WT n=25, *TAZ*$^{\Delta 45}$ n=26, WT-transfected n=9, *TAZ*$^{\Delta 45}$ n=8. Significant differences are indicated; * ≤ 0.05, ** ≤ 0.005, *** ≤ 0.0005, **** ≤ 0.00005.

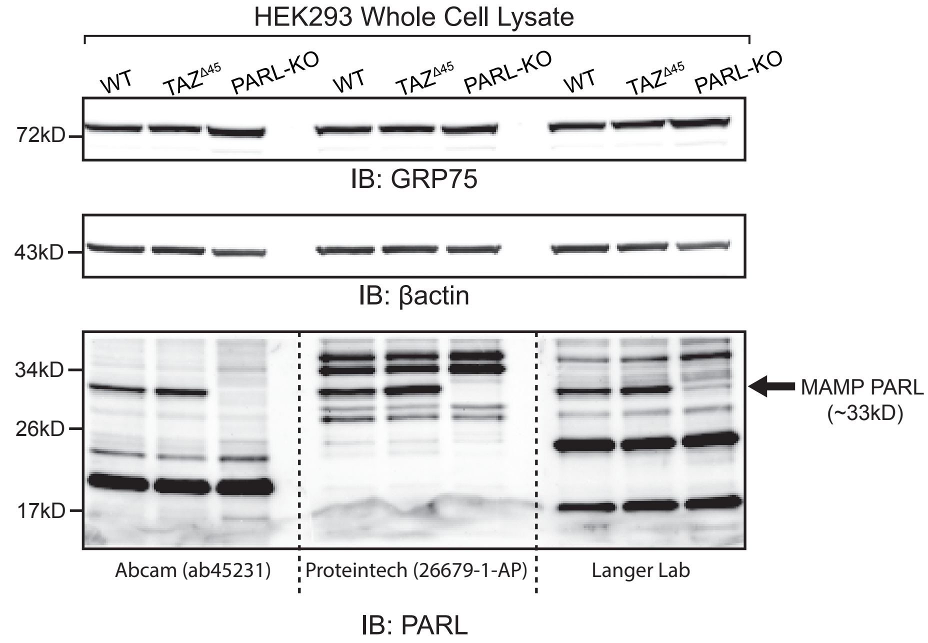


**Figure S9. Validation and classification of PARL antibodies.** Whole cell lysate (40 ug) of the indicated lines were immunoblotted for the indicated proteins. A single band at ~33kD is present in both WT and $TAZ^{\Delta 45}$ cells and absent in the PARL-KO cells, we classified this band as PARL.

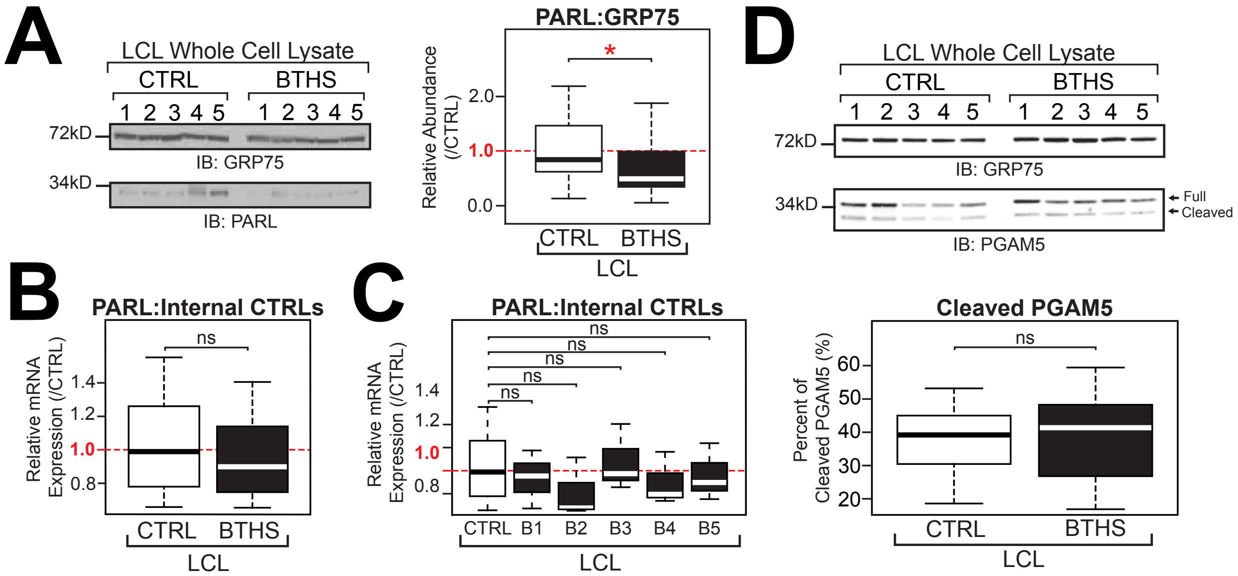


**Figure S10. PARL abundance and activity in control and BTHS LCL lines. (A)** Whole cell lysate (45 ug) of the indicated lines were immunoblotted for the indicated proteins. Band intensities, relative to loading control GRP75, were quantified and plotted relative to WT/CTRL abundance; CTRL n=26, BTHS n=35. **(B)** Relative mRNA expression of *PARL* determined by qRT-PCR and $\Delta\Delta C_T$ quantification; CTRL n=10, BTHS n=15. **(C)** Relative mRNA expression of *PARL* varies across different LCL lines; WT n=3, $TAZ^{\Delta 45}$ n=3, CTRL n=10, B1 n=3, B2 n=3, B3 n=3, B4 n=3, B5 n=3. **(D)** Whole cell lysate (45 ug) of the indicated lines were immunoblotted for the indicated proteins. Band intensities, relative to the loading control GRP75, for both full-length and cleaved PGAM5 were individually quantified and plotted as the percent of cleaved PGAM5 (cleaved/full+cleaved); CTRL n=20, BTHS n=18. Significant differences are indicated; * ≤ 0.05, ** ≤ 0.005, *** ≤ 0.0005, **** ≤ 0.00005.

**Table S4. Percent of PGAM5 cleavage with CCCP treatment (20uM) at serial time points (Figure 3D).**

| Cell Line | Time Point | n# | Percent (%) Cleaved PGAM5 | WT vs. $\mathit{TAZ}^{\Delta 45}$ For each time point | Difference (%) in Percent Cleaved |
|---|---|---|---|---|---|
| WT | 0 MIN | 41 | 12 | $p = 1.5 \times 10^{-7}$ | 11 |
| $\mathit{TAZ}^{\Delta 45}$ | | 41 | 23 | | |
| WT | 10 MIN | 8 | 11 | ns | 9 |
| $\mathit{TAZ}^{\Delta 45}$ | | 8 | 20 | | |
| WT | 30 MIN | 8 | 18 | $p = 0.001$ | 15 |
| $\mathit{TAZ}^{\Delta 45}$ | | 7 | 33 | | |
| WT | 60 MIN | 8 | 29 | $p = 3.0 \times 10^{-5}$ | 16 |
| $\mathit{TAZ}^{\Delta 45}$ | | 5 | 45 | | |
| WT | 90 MIN | 5 | 39 | $p = 2.4 \times 10^{-4}$ | 18 |
| $\mathit{TAZ}^{\Delta 45}$ | | 7 | 57 | | |
| WT | 120 MIN | 7 | 49 | $p = 7.5 \times 10^{-5}$ | 18 |
| $\mathit{TAZ}^{\Delta 45}$ | | 7 | 67 | | |

**Table S5. Abundance of PARL with CCCP treatment (20uM) at serial time points (Figure 3E).**

| Cell Line | Time Point | n# | Relative Abundance (/WT 0 mins) | WT vs. *TAZ*$^{\Delta45}$ For each time point |
|---|---|---|---|---|
| WT | 0 MIN | 54 | 1.00 | p= 1.8 x $10^{-10}$ |
| *TAZ*$^{\Delta45}$ | | 48 | 1.51 | |
| WT | 10 MIN | 5 | 1.13 | p= 1.3 x $10^{-3}$ |
| *TAZ*$^{\Delta45}$ | | 5 | 1.48 | |
| WT | 30 MIN | 6 | 1.11 | p= 2.5 x $10^{-3}$ |
| *TAZ*$^{\Delta45}$ | | 5 | 1.44 | |
| WT | 60 MIN | 6 | 1.14 | p= 6.4 x $10^{-3}$ |
| *TAZ*$^{\Delta45}$ | | 5 | 1.44 | |
| WT | 90 MIN | 5 | 0.99 | p= 5.7 x $10^{-5}$ |
| *TAZ*$^{\Delta45}$ | | 5 | 1.38 | |
| WT | 120 MIN | 6 | 1.14 | ns |
| *TAZ*$^{\Delta45}$ | | 6 | 1.42 | |

**Table S6. Relative abundance of NDUFAF1 and relevant statistics of HEK293 WT and *TAZ*$^{\Delta 45}$ cells treated with BEL and SS-31 (Figure 4A)**

<table>
<tr><th>Line & Treatment</th><th>n</th><th>Relative Abundance (/WT)</th><th>WT vs. TAZ$^{\Delta 45}$ For each treatment</th><th>TAZ$^{\Delta 45}$ vs. TAZ$^{\Delta 45}$-BEL</th><th>TAZ$^{\Delta 45}$ vs. TAZ$^{\Delta 45}$-SS-31</th></tr>
<tr><td>WT</td><td>27</td><td>1</td><td rowspan="2">p= 4.9 x $10^{-10}$</td><td rowspan="6">ns</td><td rowspan="6">p= 2.8 x $10^{-5}$</td></tr>
<tr><td>TAZ$^{\Delta 45}$</td><td>26</td><td>0.69</td></tr>
<tr><td>WT-BEL</td><td>9</td><td>1</td><td rowspan="2">p= 4.5 x $10^{-3}$</td></tr>
<tr><td>TAZ$^{\Delta 45}$-BEL</td><td>9</td><td>0.78</td></tr>
<tr><td>WT-SS-31</td><td>9</td><td>1</td><td rowspan="2">ns</td></tr>
<tr><td>TAZ$^{\Delta 45}$-SS-31</td><td>9</td><td>0.92</td></tr>
</table>

**Table S7. Relative mRNA expression and relevant statistics of HEK293 WT and *TAZ*$^{\Delta 45}$ cells treated with BEL and SS-31**

| Line & Treatment | n | Average Relative Expression to WT ($2^{-\Delta\Delta CT}$) | p-Value | | |
|---|---|---|---|---|---|
| | | | WT vs. *TAZ*$^{\Delta 45}$ For each treatment | *TAZ*$^{\Delta 45}$ vs. *TAZ*$^{\Delta 45}$-BEL | *TAZ*$^{\Delta 45}$ vs. *TAZ*$^{\Delta 45}$-SS-31 |
| **NDUFB2 (Figure S12A)** | | | | | |
| WT | 6 | 1 | p= 0.04 | ns | ns |
| *TAZ*$^{\Delta 45}$ | 6 | 0.85 | | | |
| WT-BEL | 3 | 1 | ns | | |
| *TAZ*$^{\Delta 45}$-BEL | 3 | 1.03 | | | |
| WT-SS-31 | 3 | 1 | ns | | |
| *TAZ*$^{\Delta 45}$-SS-31 | 3 | 0.94 | | | |
| **NDUFAB1 (Figure S12B)** | | | | | |
| WT | 6 | 1 | p=0.001 | p= 0.02 | ns |
| *TAZ*$^{\Delta 45}$ | 6 | 0.71 | | | |
| WT-BEL | 3 | 1 | ns | | |
| *TAZ*$^{\Delta 45}$-BEL | 3 | 0.98 | | | |
| WT-SS-31 | 3 | 1 | p=0.01 | | |
| *TAZ*$^{\Delta 45}$-SS-31 | 3 | 0.88 | | | |
| **NDUFB4 (Figure S12C)** | | | | | |
| WT | 4 | 1 | p= 0.02 | ns | p= 0.03 |
| *TAZ*$^{\Delta 45}$ | 6 | 0.71 | | | |
| WT-BEL | 3 | 1 | ns | | |
| *TAZ*$^{\Delta 45}$-BEL | 3 | 0.97 | | | |
| WT-SS-31 | 3 | 1 | ns | | |
| *TAZ*$^{\Delta 45}$-SS-31 | 3 | 1.09 | | | |
| **NDUFAF1 (Figure 4B)** | | | | | |
| WT | 6 | 1 | p= 6.4 x $10^{-4}$ | p= 0.05 | p= 0.01 |
| *TAZ*$^{\Delta 45}$ | 6 | 0.80 | | | |
| WT-BEL | 3 | 1 | ns | | |
| *TAZ*$^{\Delta 45}$-BEL | 3 | 0.94 | | | |
| WT-SS-31 | 3 | 1 | ns | | |
| *TAZ*$^{\Delta 45}$-SS-31 | 3 | 0.97 | | | |
| **NDUFB6 (Figure S12D)** | | | | | |
| WT | 6 | 1 | p= 0.01 | p= 0.05 | ns |
| *TAZ*$^{\Delta 45}$ | 6 | 0.75 | | | |
| WT-BEL | 3 | 1 | ns | | |
| *TAZ*$^{\Delta 45}$-BEL | 3 | 1.04 | | | |
| WT-SS-31 | 3 | 1 | p= 0.001 | | |
| *TAZ*$^{\Delta 45}$-SS-31 | 3 | 0.82 | | | |

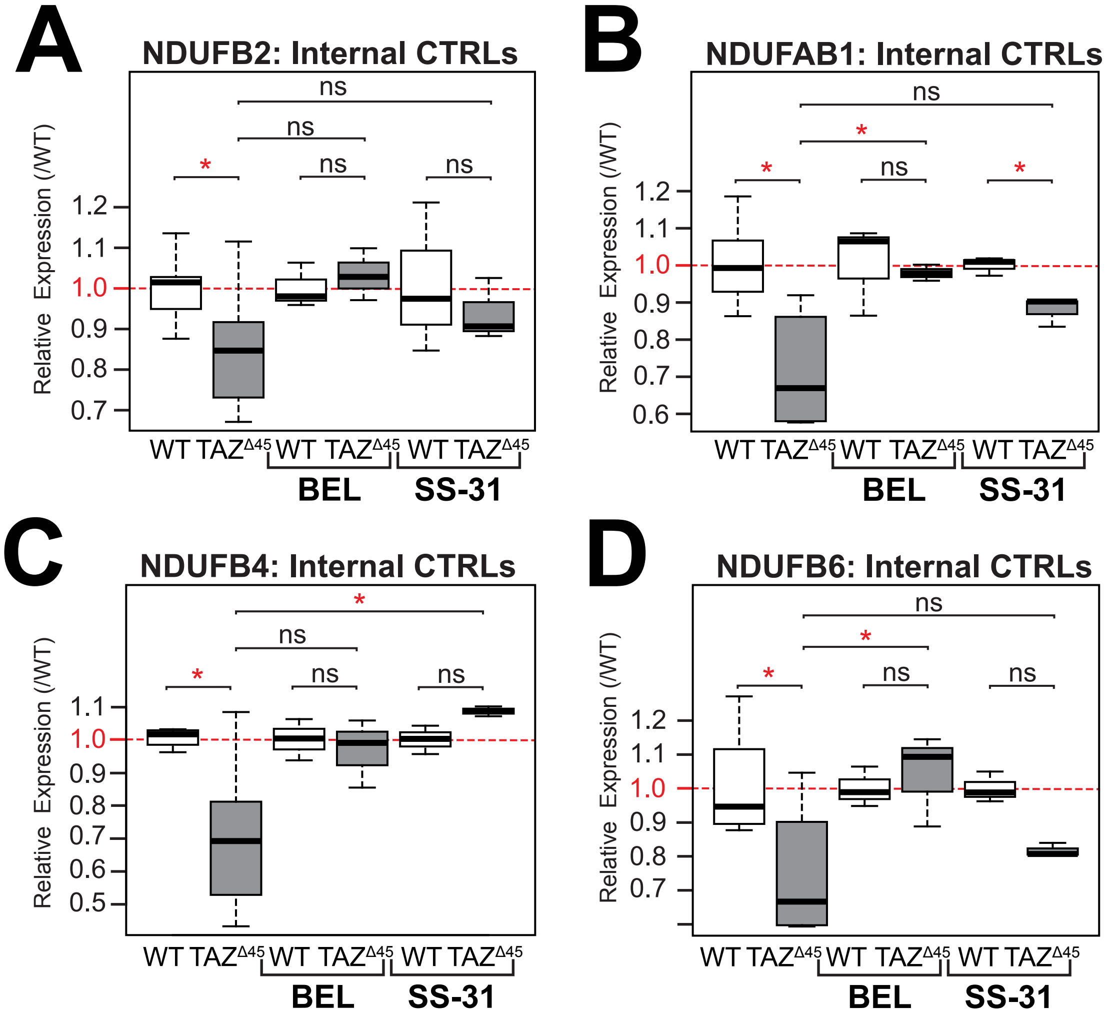


**Figure S12. Relative mRNA expression of (A) *NDUFB2* (B) *NDUFAB1* (C) *NDUFB4* and (D) *NDUFB6* after treatment with BEL and SS-31.** Expression determined by qRT-PCR and $\Delta\Delta C_T$ quantification using each respective control; WT n=6, *TAZ*$^{\Delta 45}$ n=3, WT-BEL n=3, *TAZ*$^{\Delta 45}$-BEL n=3, WT-SS-31 n=3, *TAZ*$^{\Delta 45}$-SS-31 n=3 per gene. Significant differences are indicated; * ≤ 0.05, ** ≤ 0.005, *** ≤ 0.0005, **** ≤ 0.00005.

**Table S8. CI holoenzyme abundance and relevant statistics of HEK293 WT and *TAZ*$^{\Delta 45}$ cells treated with BEL and SS-31**

| Line & Treatment | n | Relative Abundance (/WT) | WT vs. *TAZ*$^{\Delta 45}$ For each treatment | *TAZ*$^{\Delta 45}$ vs. *TAZ*$^{\Delta 45}$-BEL | *TAZ*$^{\Delta 45}$ vs. *TAZ*$^{\Delta 45}$-SS-31 |
|---|---|---|---|---|---|
| **CI:CIV (Figure 4C)** | | | | | |
| WT | 11 | 1 | p= 2.3 x $10^{-5}$ | ns | ns |
| *TAZ*$^{\Delta 45}$ | 11 | 0.59 | | | |
| WT-BEL | 8 | 1 | p= 5.2 x $10^{-3}$ | | |
| *TAZ*$^{\Delta 45}$-BEL | 8 | 0.77 | | | |
| WT-SS-31 | 3 | 1 | ns | | |
| *TAZ*$^{\Delta 45}$-SS-31 | 3 | 0.80 | | | |
| **CI:CII (Figure 4C)** | | | | | |
| WT | 10 | 1 | p= 1.6 x $10^{-4}$ | p= 6.1 x $10^{-3}$ | ns |
| *TAZ*$^{\Delta 45}$ | 13 | 0.66 | | | |
| WT-BEL | 9 | 1 | p= 0.02 | | |
| *TAZ*$^{\Delta 45}$-BEL | 10 | 0.89 | | | |
| WT-SS-31 | 4 | 1 | ns | | |
| *TAZ*$^{\Delta 45}$-SS-31 | 6 | 0.83 | | | |

**Table S9. Percentage of cleaved PGAM5 and relevant statistics of HEK293 WT and *TAZ*$^{\Delta 45}$ cells treated with BEL and SS-31 (Figure 4D)**

| Line & Treatment | n | Percentage of Cleaved PGAM5 (%) | WT vs. *TAZ*$^{\Delta 45}$ For each treatment | *TAZ*$^{\Delta 45}$ vs. *TAZ*$^{\Delta 45}$-BEL | *TAZ*$^{\Delta 45}$ vs. *TAZ*$^{\Delta 45}$-SS-31 |
|---|---|---|---|---|---|
| WT | 41 | 12 | p= 1.5 x $10^{-7}$ | p= 0.01 | p= 2.6 x $10^{-4}$ |
| *TAZ*$^{\Delta 45}$ | 41 | 23 | | | |
| WT-BEL | 16 | 15 | ns | | |
| *TAZ*$^{\Delta 45}$-BEL | 16 | 18 | | | |
| WT-SS-31 | 9 | 18 | ns | | |
| *TAZ*$^{\Delta 45}$-SS-31 | 8 | 13 | | | |

**Table S10. Relative abundance of cleaved PARL and relevant statistics of HEK293 WT and *TAZ*$^{\Delta 45}$ cells treated with BEL and SS-31 (Figure 4E)**

| Line & Treatment | n | Relative Abundance of PARL (/WT) | WT vs. *TAZ*$^{\Delta 45}$ For each treatment | *TAZ*$^{\Delta 45}$ vs. *TAZ*$^{\Delta 45}$-BEL | *TAZ*$^{\Delta 45}$ vs. *TAZ*$^{\Delta 45}$-SS-31 |
|---|---|---|---|---|---|
| WT | 54 | 1.00 | p= 1.8 x $10^{-10}$ | p= 7.1 x $10^{-15}$ | p= 9.9 x $10^{-5}$ |
| *TAZ*$^{\Delta 45}$ | 48 | 1.51 | | | |
| WT-BEL | 11 | 0.96 | ns | | |
| *TAZ*$^{\Delta 45}$-BEL | 10 | 0.93 | | | |
| WT-SS-31 | 11 | 1.03 | ns | | |
| *TAZ*$^{\Delta 45}$-SS-31 | 12 | 1.11 | | | |

**Table S11. Cell lines.**

| Line ID in Figures | Line ID | *TAZ* Genotype |
|---|---|---|
| C1 | E1 | WT *TAZ* |
| C2 | CD09 | WT *TAZ* |
| C3 | MF17 | WT *TAZ* |
| C4 | CL114 | WT *TAZ* |
| C5 | CL156 | WT *TAZ* |
| C6 | C1-ND11043 | WT *TAZ* |
| C7 | C2-ND24284 | WT *TAZ* |
| C8 | C3-ND0438 | WT *TAZ* |
| C9 | C5-ND23427 | WT *TAZ* |
| B1 | JHUHV16_007 | c.53_54delCC |
| B2 | JHUHV16_018 | c.207C>G |
| B3 | JHUHV16_022 | c.238G>A |
| B4 | JHUHV15_033 | c.82_84_delGTG |
| B5 | JHUHV16_104 | c.171delA |
| B6 | JHUHV16_060 | c.124delC |

**Table S12. Primers used for qRT-PCR.**

| Gene | Sequence (5' – 3') | |
|---|---|---|
| *TBP* | Forward | GAGCTGTGATGTGAAGTTTCC |
| | Reverse | TCTGGGTTTGATCATTCTGTAG |
| *HPRT1* | Forward | TGAGGATTTGGAAAGGGTGT |
| | Reverse | GAGCACACAGAGGGCTACAA |
| *NDUFA9* | Forward | CGCATGGGGTCACAGGTAAT |
| | Reverse | CTCGCGTCCCATTCCAGAAA |
| *NDUFS3* | Forward | TACACAGATGAGCTGACGCC |
| | Reverse | TCCAAACATGTCCCAGATCTCC |
| *MT-ND3* | Forward | ACTACCACAACTCAACGGCT |
| | Reverse | GCGGGGGATATAGGGTCGAA |
| *NDUFB4* | Forward | CATGGGAGCTCTGTGTGGAT |
| | Reverse | TTCTTTCCTATCCCTCTCAGTTTT |
| *NDUFAB1* | Forward | GCCGCCAGTATAGCGACAT |
| | Reverse | CCAAACTGTCTAAGCCCAGGT |
| *NDUFA5* | Forward | GCGGGTGTGCTGAAGAAGA |
| | Reverse | TTCCGCTTTAACCATAGCCAG |
| *NDUFB2* | Forward | GAACTCGCTCTGGAACACCT |
| | Reverse | ACTGCTGAAGATGGTGGAGT |
| *NDUFB6* | Forward | TCCATGGGGTATACAAAAAGAG |
| | Reverse | GGAAATTCTTTCATTGGTGGA |
| *NDUFAF1* | Forward | GGCAGGAGGTCAAGATTCCTT |
| | Reverse | AGCCAAGGTGAATCCTATAGAAGAG |
| *PARL* | Forward | CGCCATGGATACAGCAGGA |
| | Reverse | CACTAGCGGCTCCCTGTTCTT |
| *MT-RNR1* | Forward | TAGAGGAGCCTGTTCTGTAATCGAT |
| | Reverse | CGACCCTTAAGTTTCATAAGGGCTA |
| *MT-CO1* | Forward | GACGTAGACACACGAGCATATTTCA |
| | Reverse | AGGACATAGTGGAAGTGAGCTACAAC |
| *MT-ATP6* | Forward | TAGCCATACACAACACTAAAGGACGA |
| | Reverse | GGGCATTTTTAATCTTAGAGCGAAA |
